# A Computational Toolkit for Designing and Analysing Repeated Binary Choice Experiments

**DOI:** 10.64898/2026.01.21.700416

**Authors:** Bernard Costa, Marcus V. C. Baldo, Carolina Feher da Silva

**Author notes:** “In the natural environment of a living being, cues, means or pathways to a goal are usually neither absolutely reliable nor absolutely wrong. In most cases there is, objectively speaking, no perfect certainty that this or that will, or will not, lead to a certain end, but only a higher or lesser degree of probability” (Brunswik, 1939).

## Abstract

Adaptive human behaviour depends on the ability to detect regularities and probabilistic structures within a noisy environment. Repeated binary choice tasks, in which individuals predict one of two possible options, have long served as a fundamental tool for investigating learning, reward processing, and decision—making under uncertainty. However, traditional analyses of these tasks often rely on coarse measures such as accuracy or mean responses, overlooking the temporal information contained in behavioural sequences.

This paper introduces a mathematical and computational framework to improve the analysis of binary choice data. First, we employ higher—order Markov chains to generate sequences with controlled probabilistic dependencies, allowing for a more subtle examination of how participants extract information and learn temporal structures. Presenting analytical methods derived from time series analysis (autocorrelation, cross—correlation, shifting probabilities and Markov reconstruction) to extract structural information from simulated data of prototypical behaviours, we demonstrate how these methods can identify distinct decisional patterns that remain hidden when using conventional approaches. Finally, we validate the proposed toolkit by applying it to empirical datasets.

By revealing subtle but meaningful features of sequential decision—making, this framework enhances the interpretive power of probabilistic learning experiments. It provides researchers with tools to more accurately describe behavioural dynamics and deepens our understanding of the cognitive processes governing adaptive decisions in both healthy and clinical populations.

## Introduction

We can conceive the flow of information streaming in our perceptual world as composed of regularly occurring events embedded in a noisy environment. Hence, adaptive behaviours taking place in a consistent but chancy world have selected us for being able to detect the probabilistic structure of recurring events, allowing us to extract regularities from their noisy surroundings and increasing the likelihood of reaching desired outcomes. Therefore, grasping the relationships between successive events and learning the temporal structures of sequences is a fundamental cognitive capacity underlying human adaptive behaviour (Marković et al., 2022; Oskarsson et al., 2009).

Many of these events, certainly relevant to our everyday lives, are binary in their essence, such as a succession of rainy and sunny days or colder and warmer seasons, the sequence of failures and successes when learning a new motor task or the ups and downs of the stock market (Oskarsson et al., 2009). Repeated binary choices are not only ubiquitous elements of decision—making behaviours throughout several phyletic levels but also offer the simplest tool, albeit a very powerful one, to assess the mechanisms of probability learning in humans and other animals. Our beliefs about future events, which are crucial for the decisions that shape our behaviours, are built upon past events occurring in sequences of varying length and complexity and may depend on a broad class of learning mechanisms, which are available early in development (Aslin et al., 1998; Aslin & Fiser, 2005; Dan et al., 2025; Johnson & Munakata, 2005).

Probability learning has a long history in psychology and cognitive sciences, since Brunswik’s and Humphreys’ seminal experiments (Brunswik, 1939; Brunswik & Herma, 1951; Humphreys, 1939), and its literature is naturally intermingled with that in closely related areas or under slightly different labels, most notably statistical learning, stochastic learning, reinforcement learning and implicit learning. Its development over almost ninety years has allowed the scrutiny of cognitive mechanisms in humans — both healthy individuals and neuropsychiatric patients — as well as in several different species of other animals (Behrend et al., 1962; Behrend & Bitterman, 1961; Bitterman, 1975; Brunswik, 1939; Brunswik & Herma, 1951; Conway & Christiansen, 2001; Dotsch et al., 2017; Edwards, 1953, 1954, 1961; Erev & Barron, 2005; Falk et al., 2012; Gaissmaier et al., 2016; Gaissmaier & Schooler, 2008; Gonzalez et al., 1967; Humphreys, 1939; Myers, 1976; Tversky & Edwards, 1966; Vulkan, 2000; Worthy & Maddox, 2014).

Research employing binary choice tasks has yielded crucial insights into the mechanisms of learning, reward processing, and strategy formation in typical human cognition. For example, binary prediction tasks — such as predicting which of two lights will illuminate — have long been used in studies to uncover common heuristics and cognitive biases, such as “probability matching”, a suboptimal strategy in which individuals seem to match their choice proportions to the probabilities of the events rather than consistently choosing the more likely option (Acerbi et al., 2014; Barlow et al., 2025; Behrend & Bitterman, 1961; Edwards, 1956, 1961; Feher da Silva & Baldo, 2012; Gaissmaier & Schooler, 2008; Graf et al., 1964; Koehler & James, 2010, 2010; Montag, 2021; Newell et al., 2013; Otto et al., 2011; Saldana et al., 2022; Shanks et al., 2002; Unturbe & Corominas, 2007; Vulkan, 2000).

Such tasks have also been instrumental in exploring phenomena like the “gambler’s fallacy”, the erroneous belief that a longer run of one event increases the likelihood of the other (Burns & Corpus, 2004; Fischer & Savranevski, 2015; Hindmoor, 2006; Jessup & O’Doherty, 2011; Pelster, 2020; Roney & Sansone, 2015; Studer et al., 2015; Sun & Wang, 2010, 2012, 2012), and the debated “hot hand fallacy”, the belief that success becomes more likely for an individual after a string of prior successes (Avugos et al., 2013; Blanchard et al., 2014; Burns, 2004; Burns & Corpus, 2004; Gilovich et al., 1985; Oskarsson et al., 2009; Pelster, 2020; Stone, 2012; Sun & Wang, 2010, 2012). Furthermore, these tasks have proven invaluable in characterising altered decision—making processes associated with various neurological and psychiatric conditions, such as Parkinson’s or Huntington’s disease, often revealing altered responses to reward and risk, potentially linked to fronto—striatal pathway dysfunction (Cools et al., 2007; Lawrence et al., 1999; Morris et al., 2022; Swainson et al., 2000). Similarly, research with individuals with schizophrenia has employed such tasks to demonstrate impairments in using reward history to guide behaviour and difficulties in adapting decision strategies (Culbreth et al., 2016; Dong et al., 2024; Kasanova et al., 2011; Waltz et al., 2013a; Waltz & Gold, 2007). Studies on addiction have also utilised binary choices, often in the context of delay discounting tasks, to show how individuals with substance use disorders may devalue future rewards more steeply compared to immediate ones (Strickland et al., 2016; Vaidya et al., 2012). These examples underscore the power of binary choice tasks to provide windows into the cognitive underpinnings of both typical and pathological decision—making.

The examples above can be categorised into two broader classes of contrasting decision—making behaviours, *exploitation* and *exploration,* reflecting the well—known trade—off between them. In the former, agents tend to stay with the limited set of options that have previously yielded rewards (exploiting known opportunities), whereas in the latter, they shift their choices across a broader range of available options (exploring for potentially better opportunities). The cognitive factors involved in balancing exploitation and exploration — a challenge faced by living agents across the entire phylogenetic scale as well as throughout the development of human cognition (Cohen et al., 2007; Hills et al., 2015; Iwasaki et al., 2024; Kozunova et al., 2022; Reitich—Stolero et al., 2025; Thoma & Schulze, 2025) — can also be effectively examined through repeated binary choice tasks.

In the vast majority of reported works employing repeated binary choice tasks, the treatment of empirical binary sequences has been mostly restricted to the analysis of response proportions or accuracy measures (Barlow et al., 2025; den Ouden et al., 2013; Fantino & Esfandiari, 2002; Gaissmaier et al., 2016; Gaissmaier & Schooler, 2008; Meder et al., 2016; Miletić et al., 2021; Saldana et al., 2022; Schulze et al., 2015; Shanks et al., 2002; Shirasuna & Honda, 2023; Taylor et al., 2012; Thoma & Schulze, 2025; Unturbe & Corominas, 2007). However, an empirical binary sequence, even a short one composed of about one hundred responses, usually carries more complex temporal patterns that are not revealed by these two measures alone. Mean response and accuracy are especially limited because they become highly correlated, and therefore largely redundant, when the events of the sequence are independent and identically distributed, which is the most common experimental arrangement (this correlation does not necessarily hold, however, when the binary sequence exhibits a more structured temporal pattern). Fortunately, this valuable but frequently ignored structural content can still be accessed using a set of standard mathematical techniques, most of them developed in the context of time series analysis. To further advance our understanding of the complex cognitive processes underlying such decisions, we need mathematical tools that enhance researchers’ capabilities to understand experimental data, uncover subtle but important patterns in choice behaviour, rigorously test competing theoretical hypotheses (e.g., distinguishing between different underlying strategies that might lead to similar overall choice proportions), and ultimately gain a more nuanced understanding of human decision—making in both health and disease.

This paper intends to contribute to this ongoing effort. In Analytical Methods, we begin by introducing the use of higher—order Markov chains as input sequences (the sequences to be predicted by an observer), which capture more complex probabilistic structures than simple independent (Bernoulli) trials to generate the sequences presented to participants. Analytical Methods also introduces a range of mathematical methods which are applicable to data collected during binary choice tasks, where the “success” and “failure” outcomes are mutually exclusive and determined probabilistically on each trial (also often termed *probabilistic selection tasks* or *probabilistic learning tasks*). This distinguishes them from analyses suited for other paradigms, such as the well—known two—armed bandit task, in which each option (or “arm”) has an independent probability of yielding a gain or a loss (or neither). Following the presentation of the set of analytical tools, Description of prototypical behaviours will describe some important decisional patterns commonly found in empirical settings, as represented by prototypical virtual agents, whose stereotyped behaviours will be simulated and Simulation of Prototypical Behaviours — Results and Partial Discussion. Finally, Application of the Analytical Tools to Empirical Data Application of the Analytical Tools to Empirical Datawill demonstrate, as a validation of our analytical toolkit, its applicability to a well—established experimental paradigm known to produce predictable effects on decision—making: a binary choice task examining the differential impact of gains versus losses as payoffs on the decisional behaviour. This phenomenon, where individuals often show different learning speeds or choice patterns depending on whether outcomes are framed as exclusive gains, exclusive losses, or both in the same task (Bereby—Meyer & Erev, 1998; Erev et al., 1999; Myers et al., 1961, 1963; Myers & Suydam, 1964). The primary objective of this final step was to assess the efficacy of our proposed techniques in extracting meaningful structural elements from a real sequence of binary choices, thereby demonstrating their potential for uncovering key aspects of decision—making strategies that simpler analyses might miss. Therefore, it was intended neither as a confirmatory nor as an exploratory investigation of the empirical phenomenon; its statistical analyses were conducted solely to illustrate the implementation, interpretation, and practical utility of the analytical toolkit on a real experimental dataset.

## Analytical Methods

In a binary choice task, the participant must predict on every trial which element from a binary set will be presented next. Depending on the specifics of the experimental design, correct predictions are rewarded, incorrect predictions are punished, or both. The binary set can be any pair of contrasting values in some dimension of the perceptual space, for example, spatial location, shape, colour or semantic category of a visual stimulus (when assessing visually—mediated sensory or cognitive processes), or contrasting stimuli even in other sensory modalities, for instance in auditory or haptic dimensions (Conway & Christiansen, 2005, 2006; Mitchel & Weiss, 2011). In this section, we will offer a presentation of the methods employed (i) to generate a sequence of binary elements to be predicted (henceforth, *input* sequence) and (ii) to analyse the sequence of the observer’s predictions (henceforth, *output* sequence).

In laboratory settings, repeated binary choice tasks mediated by visual stimuli are typically conducted with participants seated in front of a computer screen on which a sequence of binary elements is displayed (see Application of the Analytical Tools to Empirical Data). On each trial, participants are instructed to predict the forthcoming binary element of the sequence, most commonly by pressing a key on the computer keyboard. Immediately following the participant’s choice — hereafter referred to as the *output* — the correct element of the sequence may or may not be presented, depending on the experimental design; when presented, this feedback determines whether the response constitutes a success or a failure.

For clarity, throughout the present text we refer to the sequence to be predicted as the *input* sequence, even though individual input elements may be revealed on each trial only after the participant’s response. Conversely, the sequence of participants’ choices is referred to as the *output* sequence. This nomenclature reflects the assumption that the temporal structure of the input sequence can shape the structure of the output sequence, whereas the reverse does not hold.

### 1. Generation of the input sequences

#### a. Markov chains

A Markov chain is a mathematical model that describes a system that moves between different possible states step by step, where the probabilities of moving to any of the next states depend only on the current state, not on the history of past states (Tomé & Oliveira, 2015).

In formal terms, for a stochastic process {X_t_}, if the probability *P*(X_t_ = n_t_) of the variable X_t_ being the state n_t_ at time t depends only on the state at the immediately preceding time step, X_t—1_, then the process satisfies the Markov property, formalised as *P*(X_t_ = n_t_ | X_0_ = n_0_,…, X_t—1_ = n_t—1_) = *P*(X_t_ = n_t_ | X_t—1_ = n_t—1_), where the notation *P*(A|B) means the conditional probability of A occurring given that B has definitely occurred.

#### b. Transition probability matrix

A transition probability matrix is a square matrix that describes all the probabilities of moving from one state of a Markov chain to another in a single time step. The entries of the matrix **T** must be nonnegative, and the sum of the entries of any row must be equal to 1.

The simplest example of a Markov chain is a sequence involving two states, labelled +1 and −1. One possible realisation of this process is the sequence +*1*−1−1+*1*−1+*1*−1+*1*+*1*−1−1. Table I below presents transition probabilities: each entry (p_1_, p_2_, q_1_, and q_2_, in yellow) indicates the probability of observing one of the states shown in the first row (in blue), given that it has been preceded by one of the states shown in the first column (in green):

**Table I.** Transition matrix for a Markov chain with two states, +1 and —1.

| | $+1$ | $-1$ |
| --- | --- | --- |
| $+1$ | $p_1$ | $q_1$ |
| $-1$ | $p_2$ | $q_2$ |

where the entries in each row sum to one: p_1_ + q_1_ = 1 = p_2_ + q_2_. From this table, we obtain the transition probability matrix **T**, given by:

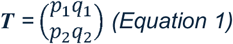

To calculate the overall frequencies of +1 and —1 in a Markov chain as well as the long—run proportion of time spent in each state, we can calculate the Markov chain’s *stationary distribution π* . The stationary distribution for a Markov chain is a two—component vector satisfying the equation:

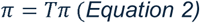

where **T** is the transition matrix and *π* is a column vector subject to the constraint that the elements of *π* are probabilities that sum to 1. Mathematically, this defines *π* as the normalised right eigenvector of **T** corresponding to the eigenvalue 1. Solving this equation and the constraint results in a linear system which can be solved manually or with tools such as *SymPy* (Meurer et al., 2017). Alternatively, the stationary distribution can be obtained by simulating a very long Markov chain and calculating the frequency of each state in this chain.

#### c. Order of the Markov chain

We can also construct a Markov chain that depends not only on the immediately preceding time step, but also on the steps prior to that. The number of previous steps, k, determines the order of the Markov chain. In particular, in a binary sequence generated by a *k*th—order Markov chain, the probability of occurrence of each possible element (—1 or +1 in the present study) at step *k*+1 is conditional on the preceding *k* elements.

For example, in a zeroth—order Markov chain, also called a Bernoulli process, the probability of each element is independent of previous elements (i.e., the state is empty); that is, *P*(X_t_ = n_t_ | X_0_ = n_0_,…, X_t—1_ = n_t—1_) = *P*(X_t_ = n_t_ ). In this case, the only thing to be learned is which element is more frequent than the other. In the case where *P*(−1) = *P*(+1) = 0.5, the sequence is completely random and unpredictable.

In a first—order Markov chain, already described above as the default definition of a Markov chain, the states correspond to one element. This allows for the generation of sequences whose elements can be “sticky” (e.g., +1 tends to be followed by +1) or “alternating” (e.g., +1 tends to be followed by −1).

In a second—order Markov chain, the state corresponds to two consecutive elements. Thus, there are 2^2^ = 4 states: +1+1, +1−1, −1+1, and −1−1. The probability of the next element depends on which of these four states has just occurred.

In the case of orders k > 1, the transition matrix, describing the probability of the next element given the previous k elements, is rectangular, not square (see Table II). However, to calculate the Markov chain’s stationary probabilities, we first need to construct an equivalent square transition matrix. This construction effectively represents the higher—order chain as a standard first—order Markov chain whose states are composed of multiple sequence elements. In this representation, transitions occur as a “sliding window” along the sequence. For example, in a second—order chain, a possible state is +1−1, which can transition to −1+1 or −1−1 depending on whether the next element is +1 or −1, respectively. In contrast, transitions from +1−1 to states +1+1 or +1−1 are not structurally admissible because the element at the second position of the current state, −1, must shift to the first position of the next state. When representing a higher—order Markov chain as a first—order Markov chain, the transition matrix is square and allows for the calculation of the stationary distribution, although the resulting matrix will be sparse due to the constraints on admissible transitions.

**Table II.**
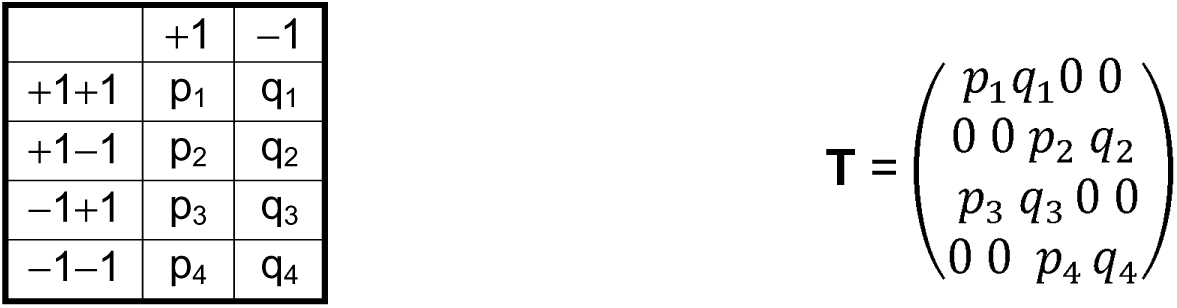
Rectangular (left) vs. square (right) transition matrices for a second—order Markov chain.

In this paper, we define Markov chains using rectangular transition matrices (see Table II). For higher—order Markov chains, this representation directly maps the history of previous elements to the probability of the next single element. By contrast, the square transition matrix is less intuitive for this purpose: it is dominated by structurally inadmissible transitions and obscures the fact that participants predict only one item per trial.

#### d. Calculating Overall Event Probabilities

Researchers often need to control the overall frequency of +1 and —1 to distinguish between learning temporal structures versus learning simple frequency biases.

For a first—order chain, the states are single elements of the sequence, −1 and +1. Hence, the stationary distribution directly gives the overall probability of each element. In a higher—order chain, the states are subsequences. To find the marginal probability of an event (e.g., +1), we simply sum the stationary probabilities of all states that end with that element.

#### e. Conditional Entropy and Sequence Predictability

Another mathematical concept that can be used to design input sequences is the conditional entropy (Cover & Thomas, 2006), which serves as a measure of the “randomness” of the input sequence from the perspective of an ideal predictor (a predictor that achieves maximum accuracy). For example, in a perfectly alternating sequence, where +1 is always followed by −1 and vice versa, while the overall probability of +1 and −1 might suggest a sequence that is highly random (50/50 split), the internal structure makes it entirely predictable.

The conditional entropy H(X|S) is the average uncertainty of the next element given the current state S, weighted by how often each state occurs (the stationary distribution π).

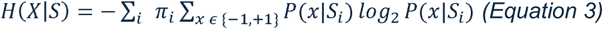

Since − log_2_ 1 = 0 , when the next element of the Markov chain can always be predicted with certainty (the probability is 1), the conditional entropy is equal to 0 bits. When the next element can be either −1 or +1 with equal probability, i.e., *P*(+1) = *P*(−1) = 0.5, since − 2 × (0.5 log_2_ 0.5) = 1 , the conditional entropy is equal to 1 bit. Hence, the conditional entropy varies from 0 (completely predictable) to 1 (completely unpredictable) bit.

This measure highlights the benefit of learning the sequence structure. A participant who treats a perfectly alternating sequence as a zeroth—order Markov chain, where the probability of both +1 and −1 is 0.5, faces the full entropy, 1 bit. A participant who successfully learns the Markov dependencies as a first—order Markov chain reduces their uncertainty to the conditional entropy of 0 bits (in this specific case; in other cases, the conditional entropy will be reduced but not necessarily to 0 bits). The difference (1 − 0 = 1 bit) represents the information gained by learning the temporal pattern.

#### f. Model Reducibility

When designing complex patterns, researchers must ensure the chain is not reducible to a lower order. A transition matrix defined for order k collapses to order k−1 if the oldest element in each state does not influence the probability of the next element, that is, if the rows of the transition matrix are the same for states with the same oldest element.

#### g. Markov chains used in this study

The present work’s input sequences were generated by two different Markov processes: a Markov chain of order zero (M0) and a Markov chain of order two (M2) (see below). Table III shows the transition matrices used in each one of these processes. In M0, the binary element x_i_ assumes the values +1 or −1 with probabilities p and 1 − p, respectively (in all analyses reported in this work, p = 0.67 and consequently 1 − p = 0.33). In M2, the probabilities of either +1 or −1 occurring depend on the two previous events. For example, when two consecutive −1s are presented, a +1 will occur next with certainty (this probability can be read from the fourth row of the M2 matrix, Table III). However, when two +1s happen consecutively, a +1 can occur next with 0.67 probability, whereas a −1 can occur next with 0.33 probability (first row in Table III).

**Table III.**
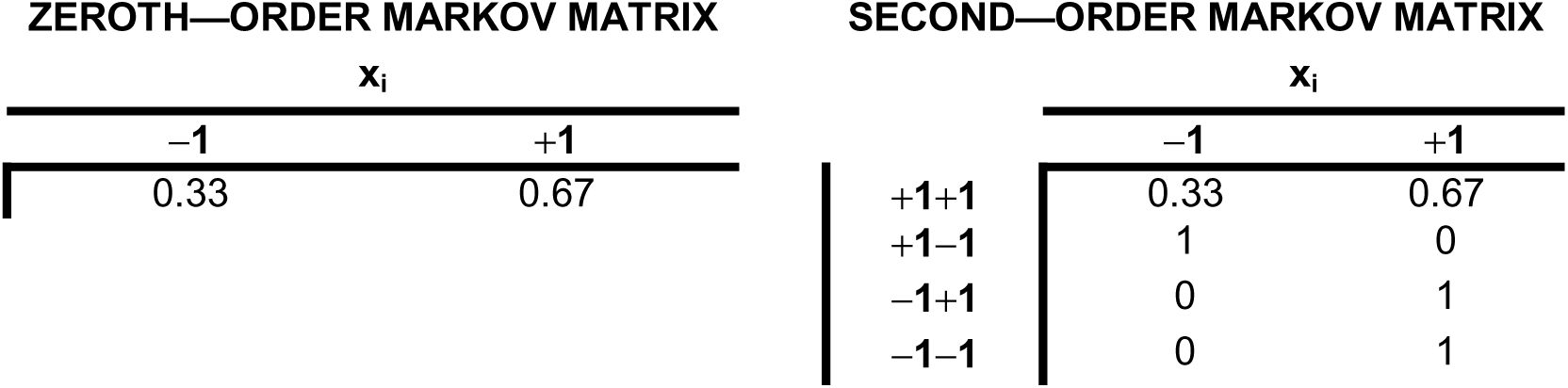
Markov transition matrices used to generate the two kinds of sequences employed throughout the present work in simulating prototypical behaviours (Simulation of Prototypical Behaviours Results and Partial Discussion) as well in running experimental procedures (Application of the Analytical Tools to Empirical Data).

Because of the choice of the elements in the transition matrices designed for the present work (Table III), both M0 and M2 sequences intentionally exhibit the same asymptotic probability distribution for the binary events +1 and −1, respectively, p = 0.67 and 1 − p = 0.33. This result is trivial for M0. For M2, it can be proven by calculating the stationary probabilities of the four states, which are approximately [0.503, 0.166, 0.166, 0.166] for the states in the same order as shown in the first column of Table III. The states ending in +1 are +1+1 and −1+1, and thus the overall (asymptotic) probability of +1 is 0.503 + 0.166 = 0.668. Therefore, the probability of −1 is 1 − 0.668 = 0.332.

The conditional entropy of the two processes can be calculated from their transition matrices and stationary probabilities. In the case of the M0 sequence, the conditional entropy is simply − 0.33 log_2_ 0.33 − 0.67 log_2_ 0.67 ≈ 0.915. Because the M2 sequence has the same asymptotic probabilities for +1 and —1 as the M0 sequence, a participant who hasn’t learned the temporal structure of the M2 sequence will face 0.915 bits of entropy for each element. However, an ideal predictor who takes into account the two previous elements to predict the next would face a conditional entropy of only 0.503 × (− 0.33 log_2_ 0.33 − 0.67 log_2_ 0.67) + 3 × 0.166 × 0 ≈ 0.46 . Therefore, learning the temporal structure of the M2 sequence nearly halves the entropy of each element.

### 2. Measures utilised in the analysis of the binary sequences

Here we will present some techniques that are appropriate to analyse discrete time series. In the following discussion, X and Y will represent the input and output binary sequences, respectively, i.e., the sequence chosen by the experimenter or generated by a computer (X), and the sequence of predictions made by an observer, real or simulated (Y). To clarify the order of events during an experiment with human participants, in each trial i, the participant makes a prediction y_i_ and then observes the input x_i_ and the achieved outcome (win or loss).

#### a. Mean response

For a binary sequence of length N, with elements coded as −1 and +1, the proportion q (or frequency) of +1s provides an estimate of the asymptotic stationary probability of observing the event coded as +1 — hereafter referred to as the “mean response” — and is given by:

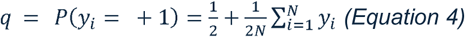

Clearly, the stationary probability of the event −1 is given by *P*(y_i_ = −1) = 1 − q, and we arbitrarily elected the frequency of +1s as a measure of mean response in the present analyses, coding the most frequent binary event in our input sequences.

#### b. Accuracy

This measure provides the proportion of correct responses (predictions) made by the participant and therefore reflects the similarity between the input and output sequences. The accuracy is a measure related to the cross—correlation between the input and response sequences at lag τ = 0 (see below), and for sequences whose elements are coded as −1s and +1s can be calculated as:

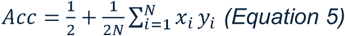

The accuracy can also be separately calculated for several disjoint, contiguous, or superimposed segments of the output sequence, or even by a moving average algorithm, thus allowing the construction of a learning curve that reflects the participant’s learning kinetics.

##### 1. Maximum Achievable Accuracy

When a binary input sequence is generated by a k—th order Markov chain, the maximum achievable accuracy *Acc_max_* is the upper limit of correct predictions an ideal predictor can make. In the present context, an ideal predictor may be conceived as an agent capable of estimating the transition probabilities of a Markov matrix, at least to the extent of determining, for each row of the transition matrix, which of the two complementary probabilities is larger.

For any state S in a k—th order Markov chain, the optimal prediction *ȳ* for the next element of the output sequence is the value (−1 or +1) with the highest conditional probability, given by the transition matrix:

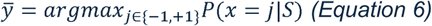

The probability of being correct in state S is simply the probability of the more likely result, max{*P*(x = −1|S), *P*(x = +1|S)}. To find the maximum accuracy across the entire sequence, we calculate the expected value of these maximum probabilities for each state, weighted by the state’s stationary probability, given by π (see b. Transition probability matrix):

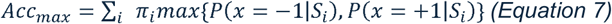

The maximum accuracy is a deterministic function of the conditional entropy: as uncertainty increases, the ceiling for performance necessarily drops. However, while the conditional entropy quantifies uncertainty in bits, the *Acc_max_* provides a direct measure of behavioural performance in terms of hit rate. A sequence with a conditional entropy of 0 bits is entirely deterministic, allowing for *Acc_max_* = 1 (perfect prediction). Conversely, a sequence with 1 bit of entropy per trial is purely stochastic, limiting the ideal predictor to *Acc_max_* = 0.5, which represents performance at chance level.

As examples, we can calculate the maximum accuracies for the M0 and M2 Markov chains used in this study (Table III). The next element in the M0 chain can be predicted based on the overall probabilities of —1 and +1, which are 0.33 and 0.67, respectively. Therefore, an ideal predictor would always predict that the next element is +1 and obtain an accuracy of 0.67. The M2 Markov chain has deterministic transitions for three of the four possible states, which lead to certain predictions, and the remaining state is followed by —1 and +1 with probabilities 0.33 and 0.67. Because the stationary probability of the latter state is approximately 0.503, the maximum accuracy is (0.503 × 0.67) + (0.497 × 1) ≈ 0.834.

#### c. Cross—correlation function

The cross—correlation is a measure of the similarity between two sequences of the same length. For two binary sequences *X=*(*x_1_ ,x_2_ ,x_3_,…,x_N_*) and *Y =* (*y_1_, y_2_, y_3_,…, y_N_*), both of length N and composed of +1s and −1s, the cross—correlation CC_X,Y_ can be defined simply as:

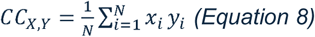

where —1 ≤ CC_X,Y_ ≤ 1 for any pair of sequences X and Y, as long as their binary elements are +1s and −1s.

The cross—correlation also allows us to calculate the similarity between a subsequence Y_s_ of the output sequence Y and a subsequence X_s_ of past (or future) elements of the input sequence X, which starts a fixed number of trials before (or after) the start of the output subsequence. For different numbers of trials (lags) τ, where τ can range from −(N − τ) to (N − τ), the cross—correlation function (CCF) C_X,Y_(τ) can be defined as:

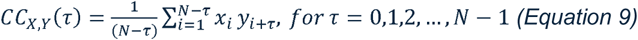

and

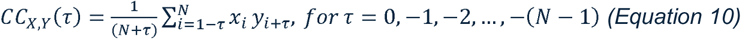

Equation 9 gives the cross—correlation coefficient, as a function of τ, between a subsequence Y_s_ of Y (the output) and a subsequence X_s_ of X (the input), where X_s_ is shifted τ trials into the past in relation to Y_s_; Equation 10, conversely, provides the cross—correlation coefficient between a subsequence Y_s_ of Y and a subsequence X_s_ of X that is shifted τ trials into the future. As long as both sequences X and Y are composed of +1s and −1s, the cross—correlation coefficients are confined to the interval −1 ≤ *CC_X_*_,*Y*_(*τ*) ≤ 1 for any X, Y, and τ; also, as can be easily verified, Equation 8 is a special case of Equation 9 or 10 for τ = 0 and relates to the accuracy (Equation 5) by the relation 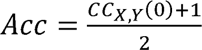. Splitting the CCF into Equations 9 and 10 allows us to examine separately the potential causal influence of past and future events in the input sequence on present events in the output sequence (Equation 9). By contrast, no such causal relationship can exist between present input events and future output events (Equation 10), which naturally leads to an asymmetry of the CCF around τ = 0. This asymmetry can, in turn, reveal the extent to which features of the input sequence affect the agent’s decisional behaviour.

This formulation of the CCF is called aperiodic, and we adopted it instead of the periodic formulation because the sequences analysed in this study are finite and non—repeating. The periodic formulation assumes that the observed sequences are circular and repeat indefinitely over time, as if their last element were followed again by their first element. Such an assumption is violated for behavioural data. In contrast, the aperiodic formulation respects the causal and temporal structure of the task by treating values outside the observed interval as undefined.

A consequence of this choice is that the number of overlapping terms used to compute the cross—correlation decreases as the lag increases, since fewer elements remain available for comparison. This reduction limits the reliability of estimates at larger lags and can make their computation unfeasible in short sequences. By contrast, the periodic formulation avoids this issue because it treats the sequence as circular, preserving a constant number of terms at all lags. Nevertheless, despite this limitation, the aperiodic formulation remains the most appropriate for the present context.

#### d. Autocorrelation function

Analogously to the cross—correlation function, we can measure the similarity of a sequence with itself, given a time lag τ. Such a measure, called the autocorrelation function (ACF), is given by:

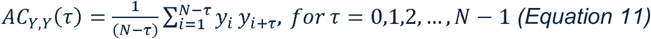

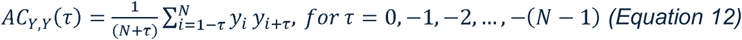

In fact, the autocorrelation function is the cross—correlation function between a sequence and itself (again, −1 ≤ AC_X,X_ ≤ 1 and —1 ≤ AC_Y,Y_ ≤ 1). For every sequence, the autocorrelation coefficient at lag τ = 0 is trivially equal to unity: AC_X,X_(0) = AC_Y,Y_ (0) = 1. For the ACF, in contrast with the CCF, Equations 11 and 12 yield the same result when τ = ±k for any admissible k, meaning that the ACF is symmetrical around τ = 0.

#### e. Shifting probabilities

We can define a measure that quantifies the participant’s tendency to switch (shift) their choice relative to the immediately preceding one, either unconditionally or conditionally on whether the previous choice was correct (“win”) or incorrect (“lose”). More formally:

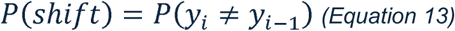

where *P*(shift) measures the unconditional probability of a given choice (y_i_) being different from the previous one (y_i—1_). The conditional shifting probabilities are given by the following equations:

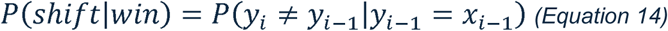

and

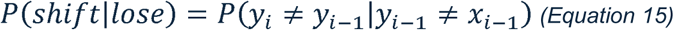

Thus, Equations 14 and 15 measure the probability of the participant choosing an option that is different from the previous one (a shift, y_i_ ≠ y_i—1_) conditionally on the previous outcome being, respectively, a “win” (y_i—1_ = x_i—1_) or a “loss” (y_i—1_ ≠ x_i—1_).

The shifting diagram (Figure 1) represents the relationship between the probability of shifting after a loss and the probability of shifting after a win and may reveal the predominance of four different decisional behaviours. For instance, if after a loss the agent shows a higher probability of shifting to the alternative option than of remaining with the currently chosen one, and after a win it shows the opposite pattern, the behaviour can be classified as “win—stay/lose—shift”. In this case, the agent’s behaviour will be represented by a point in the diagram’s upper—left quadrant. A similar interpretation applies to the other behavioural patterns, which correspond to the remaining three quadrants of the shifting diagram.

**Figure 1.**
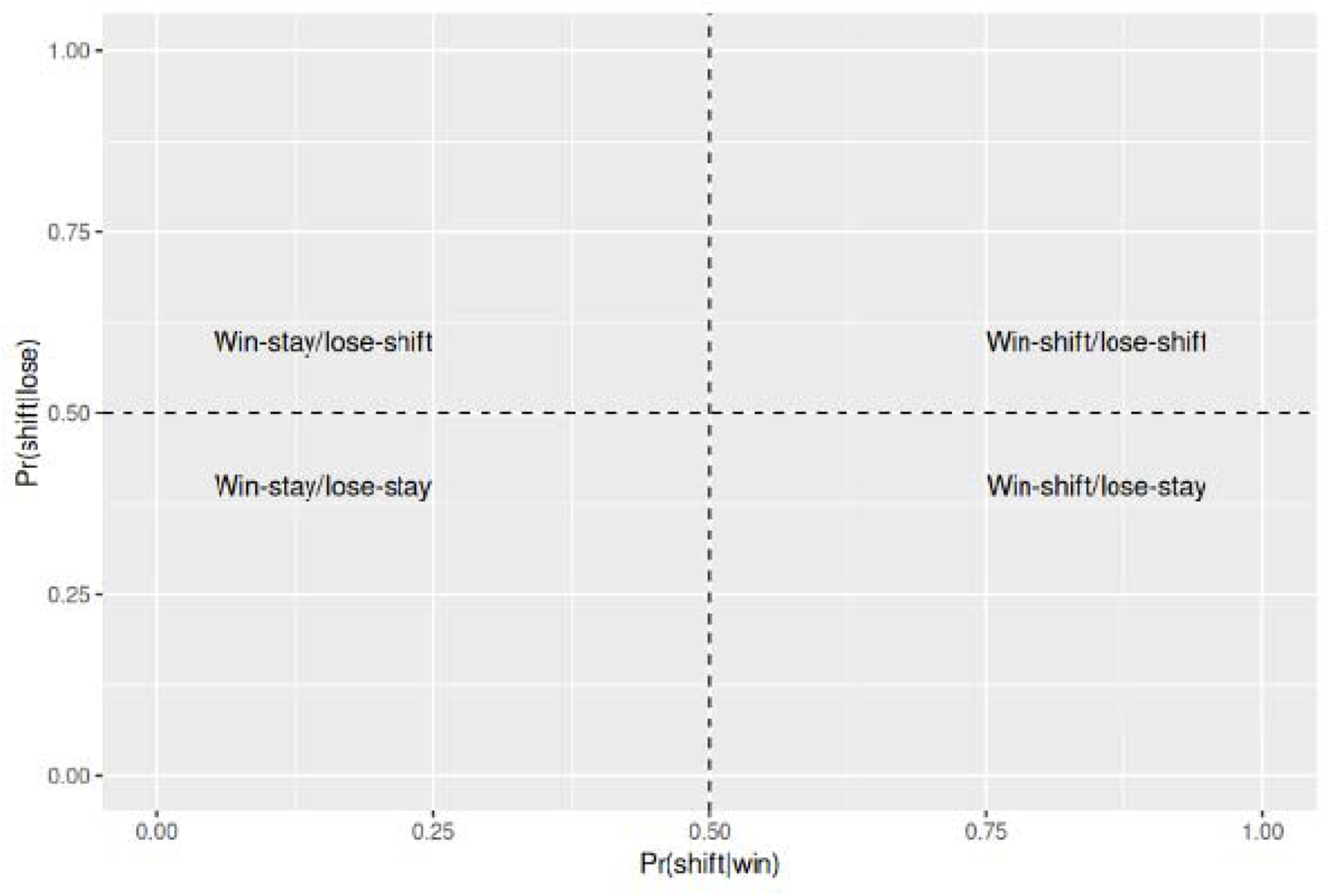
Shifting diagram showing the relationship between the probability of switching after a loss and the probability of switching after a win. The distribution of points across the four quadrants indicates different decision—making patterns, such as “win—stay/lose—shift” and its alternatives.

#### f. Reconstruction of Markov matrices

The simplest method to estimate the matrix of a putative Markov process (of a given order) that possibly generated a binary sequence is by estimating, for that specific order, the conditional probabilities of occurrence of the two possible events in the sequence (+1 or —1) depending on the history of recent events in the input sequence.

In a probability—learning task, task performance is determined by how the history of the input sequence influences each event in the output sequence rather than by how each event in the output sequence might have been influenced by its history. In order to reconstruct a putative Markovian process that may represent the stochastic relationship between the input and output sequences, we calculate the conditional probabilities that express the dependencies between each event of the output sequence on one, two, or more previous events of the input sequence, which result in first—, second— or higher—order transition matrices, respectively. This means that when reconstructing a second—order Markov process that, for example, represents the stochastic structure of an output sequence, Y, it is more meaningful to estimate *P*(y_i_|x_i—1_x_i—2_), where x_j_ belongs to the input sequence X, than *P*(y_i_|y_i—1_y_i—2_).

For a zeroth—order process, this is simply done by estimating the asymptotic probabilities of +1 and —1, since there is no serial dependence between events. For a first—order process, the following probabilities should be estimated: *P*(+1|+1), *P*(+1|−1), *P*(−1|+1) and *P*(−1|−1) but, since *P*(+1|+1) + *P*(−1|+1) = 1 and *P*(+1|−1) + *P*(−1|−1) = 1, only two estimates must be made. In the case of an N^th^—order process, 2^N^ independent elements of the Markov matrix are to be estimated, and, as N grows larger, this poses a problem for short sequences due to the increasingly scarcer samples for each matrix element. Estimating these conditional probabilities is equivalent to estimating the conditional mean response, since, for binary responses coded as +1 and −1, the conditional mean uniquely determines the corresponding conditional probabilities and therefore contains the same information about the underlying transition structure.

#### g. Data transformation

The measures discussed so far complement each other, offering a multivariate view, under different perspectives, of the same object: the output binary sequence provided by a decisional agent. In cases where these measures are to be subjected to inferential statistical analyses — particularly those requiring assumptions such as normality and homogeneity of variances — it is advisable to apply data transformations that help them meet those assumptions. The probability (or frequency) and proportion measures, which are confined to the interval [0,1], may be submitted to a normalised version of the Freeman—Tukey transformation before entering the statistical analysis; similarly, the autocorrelation and cross—correlation coefficients, which range from —1 to +1, may be submitted to Fisher’s r—to—z transformations (Quinn & Keough, 2002; Sheskin, 2011; Tabachnick & Fidell, 2013; Zar, 2009).

## Description of prototypical behaviours

To gain some insight into the meaning of the analytical techniques presented here, we will consider a small number of simple and intuitive strategies that may take part in more complex, real—life decisional behaviours. It is unlikely, however, that real binary responses will reflect a single, pure instance of any of these simplistic strategies. We will consider five simple decisional behaviours, labelled as “Perseveration”, “Alternation”, “Win—Stay/Lose—Shift”, “Probability Matching”, and “Ideal Predictor”.

Once we have defined the main features of these five simpler “toy model” behaviours — whose descriptions are based solely on observable responses, without any consideration of the underlying cognitive processes — we will proceed to the analysis of more complex real data. Note that the specific results presented below depend to a large extent on the particular input sequences used in the simulations, which were generated from two fixed Markov transition matrices (Table III). Thus, they should not be generalised to other input sequences produced by different Markovian processes.

### 1. Perseveration

In the present context, we define Perseveration as the tendency of an agent to repeat the same choice throughout the entire task, regardless of whether this behaviour is causally related to the input sequence to be predicted or to prediction accuracy (correct or incorrect responses). Because this response pattern can yield optimal performance when the input sequence is generated by a zeroth—order Markov process (i.e., a sequence of Bernoulli trials) and the agent perseverates in choosing the more frequent event, it has often been referred to as “maximisation” (Maddox & Bohil, 2004; Schulze et al., 2015; Unturbe & Corominas, 2007). We nonetheless prefer the term “perseveration” for two reasons. First, if the agent perseverates on the “wrong” choice, i.e., the least frequent element of an input sequence generated by a zeroth—order Markov chain, the resulting performance corresponds to the minimum accuracy. Second, even if the agent perseverates on the most frequent element of the input sequence, performance will not be optimal when the sequence exhibits a richer informational structure, as is the case when it is generated by a higher—order Markov process.

### 2. Alternation

In this simple response pattern, the participant continuously alternates between the two available options, regardless of whether this has any causal connection with the input sequence. This behaviour has been observed in kindergarten—aged children (Derks & Paclisanu, 1967).

### 3. Probability Matching

As already described in the Introduction, we will adopt this label to define the behaviour in which the frequency of the binary events in the output sequence nearly matches the frequency in the input sequence. Importantly, responses are generated independently across trials, with no temporal structure or dependence on previous events.

### 4. Win—Stay/Lose—Shift

The extent to which a behaviour conforms to a Win—Stay/Lose—Shift (WSLS) strategy is defined by the probability of repeating the previous choice when it was correct, *P*(stay|win), and the probability of shifting to the other alternative when the previous choice was incorrect, *P*(shift|lose). In a pure WSLS strategy, *P*(stay|win) = 1 = *P*(shift|lose). Note that this behaviour is empirically indistinguishable from simply repeating the immediately preceding element of the input sequence, without taking into account whether the previous outcome was a win or a loss (Cohen et al., 2007; Westerman et al., 2025).

### 5. Ideal Predictor

This behaviour is defined as the one that maximises accuracy by optimally utilising all the available statistical information about the task. Therefore, the Ideal Predictor will be tailored to the specific input sequence presented to the agent. When predicting an input sequence generated by a zeroth—order Markov chain, the Ideal Predictor consists of an agent whose response is always the most frequent event in the input sequence (in this — and only this — scenario, the behaviour of the Ideal Predictor coincides with that of a Perseverator). As for the prediction of a kth—order Markov chain, the Ideal Predictor characterises an agent that always selects the most likely option given the k previous elements of the input sequence, thereby maximising prediction accuracy. In practical terms, this implies that reconstructing the Ideal Predictor’s Markov transition matrix from an output sequence generated by such an agent would not recover the original Markov matrix that generated the corresponding input sequence. Instead, it would yield a matrix composed solely of 0s and 1s, in which the largest element in each row of the original transition matrix is replaced by 1 and the complementary element by 0.

## Simulation of Prototypical Behaviours — Results and Partial Discussion

For each one of the five behaviours defined above, we simulated virtual agents performing a repeated binary choice task where the input sequences were generated by zeroth— or second—order Markov chains (see Table III). Each agent performed a distinct binary choice task of 220 trials. The first 20 trials were always discarded, and only the remaining 200 trials were submitted to numerical analyses so that both the input and the output sequences could be considered approximately stationary.

For each Markov transition matrix, M0 and M2, 50 realisations of each prototypical behaviour were performed. For all M0 and M2 input sequences, the proportion of +1 was kept in the interval [0.64, 0.70] to ensure that the event frequencies would be similar for all agents. To introduce some degree of noise into the simulations, the probability of a given behaviour was set to 95%. For example, in the Alternation behaviour, an agent alternated from one choice to another with 95% probability and repeated the previous choice with 5% probability. The same amount of noise was introduced into the Perseveration, Win—Stay/Lose—Shift and Ideal Predictor behaviours. Since, in the Probability Matching behaviour, the agent randomly chooses the two binary events with a probability that matches the respective frequency of the binary events in the input sequence, no extra noise was added. Although computational power did not impose any relevant constraints on the simulations, a relatively small sample size (50 realisations for each type of prototypical behaviour and input sequence) and sequence length (200 trials) were chosen to reflect conditions commonly encountered in real experimental settings.

Besides being submitted to the numerical analyses, every output sequence, Y, was used to construct a set of 100 “surrogate” sequences generated by random permutation of the elements in the original sequence (Prichard & Theiler, 1994; Theiler et al., 1992). This caused the frequencies of binary events within each surrogate sequence to remain the same as for the original data set, but all serial correlations were destroyed, including any serial cross—correlation between the original output sequences Y and their respective input sequences X. The generation of surrogate data allows us to define baseline confidence intervals for summary statistics assuming no serial correlations in the original data — the null hypothesis. Using the original data to construct a null hypothesis distribution allows for statistical hypothesis testing in situations where traditional parametric methods are unsuitable or inapplicable (Prichard & Theiler, 1994; Theiler et al., 1992). In empirical scenarios, if the 95% confidence interval of a statistic estimated from a time series does not encompass the corresponding value obtained from its surrogate, one may reject the null hypothesis that the series lacks temporal dependence among its elements.

### 1. Mean response

Figure 2 shows the mean response (frequency of +1s) for the five prototypical behaviours for the two orders (M0 and M2) of Markov chains (teal lines). For the sake of comparison, the figure also shows the frequency of +1s for the set of input sequences (red lines), which are trivially the same for the five behaviours. As expected, the input sequences show a mean response in agreement with the expected frequency of the event +1 (around 0.67) for both orders of the Markov sequences and all five behaviours. Small deviations are due to the analysis being run only on 200 elements of the input sequence. Greater variability was found in the simulated output sequences, in agreement with the predominant behaviour at hand. In those behaviours that preserve the probability distribution of the input sequence, either by replicating it (WSLS) or by matching its frequencies (Probability Matching), no relevant differences were found in the mean responses between the input and output sequences. For the Alternation and Perseveration behaviours, the output mean response simply reflects their respective predominant strategies (approaching 50% in the former and 100% in the latter). As for the Ideal Predictor strategy, in a zeroth—order Markov sequence, it exhibits the same behavioural pattern as the Perseveration strategy. In a second—order Markov sequence, it maximises the probability of selecting the most likely option conditioned on the two previous events. Two aspects should be noted here: (i) an Ideal Predictor will not in general reach a mean response of 100% when the sequence to be predicted exhibits some serial correlation; (ii) for Markov chains of order 1 or greater, the Ideal Predictor’s mean response and accuracy are not necessarily correlated.

**Figure 2.**
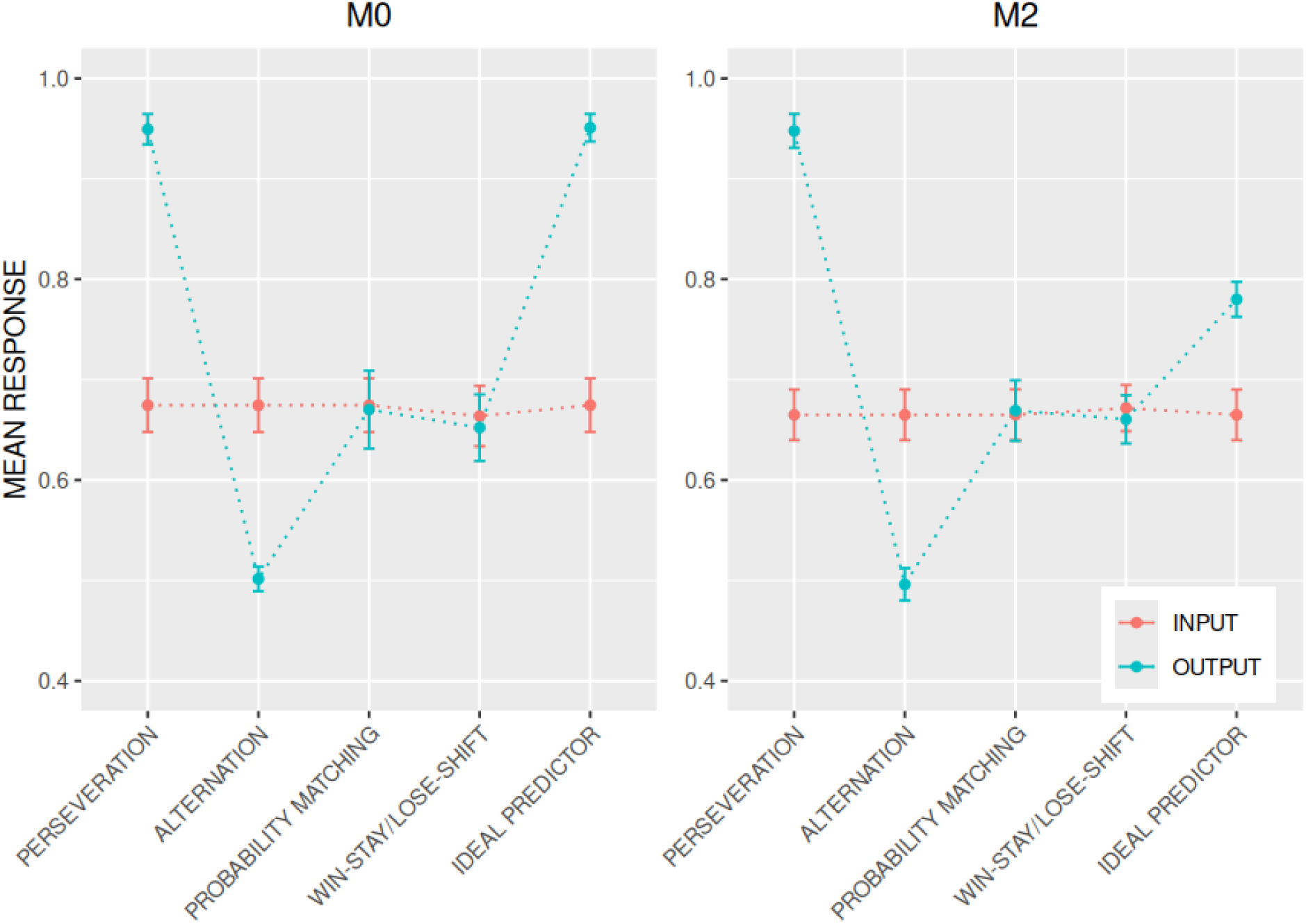
Mean response (frequency of +1s) averaged over 50 output sequences for each behaviour in each group, expressing five prototypical behaviours for both kinds of input sequences, M0 (left) and M2 (right): Alternation, Win—Stay/Lose—Shift (WSLS), Perseveration, Probability Matching, and Ideal Predictor (teal lines). Each agent performed two repeated binary tasks, one for each Markov order (M0 and M2). The frequency of +1s in the input sequences is also shown (red lines) for the sake of comparison. Error bars are ± 1 standard deviation. Surrogates are not shown, as their statistical distributions are trivially identical to the distributions of the output sequences.

### 2. Accuracy

Figure 3 shows the accuracy of the virtual agents’ responses (in red) as well as that of the surrogate sequences (in teal). These results are easily understood in relation to their respective behaviour. In Alternation, a mean response of 0.5 irrespective of the input sequence naturally leads to an accuracy of about 0.5; an accuracy of about 0.67 is expected in Perseveration, since the agent chooses the most frequent event (+1) in 95% of the trials; in Probability Matching, the expected accuracy is (0.67)^2^ + (0.33)^2^ ≅ 0.56, very close to the observed accuracy. For WSLS in the M0 condition, since the input sequence is replicated by the agent apart from 5% noise, the result is very similar to the one observed in Probability Matching: two random sequences (input and output) with the same probability distributions, therefore leading to an accuracy close to 0.56. For WSLS in the M2 condition, the tendency to replicate the input sequence produces an output that happens to align closely with the structure of the M2 sequence. This explains the higher accuracy observed in WSLS for the M2 task, inferior only to the accuracies of the Ideal Predictor and Perseveration behaviours. Crucially, this only occurs because the specific transition matrix used to generate the M2 sequences in this study incidentally favours a WSLS pattern. If, for example, we had used a different transition matrix that does not favour a win—stay/lose—shift pattern, the WSLS behaviour would have achieved a lower accuracy in the M2 task compared to the M0 task.

**Figure 3.**
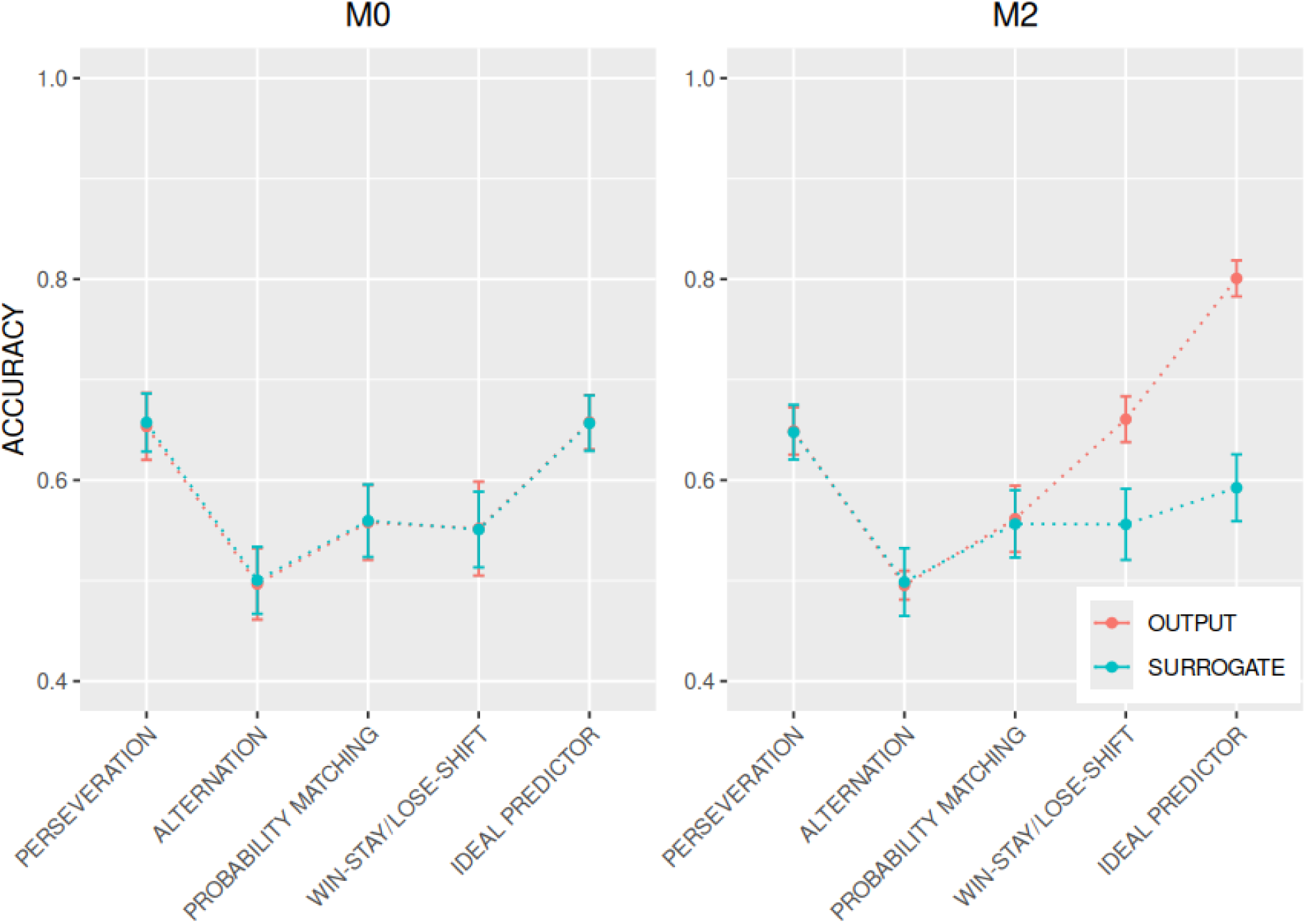
Accuracy averaged over 50 output sequences in each group expressing five prototypical behaviours: Alternation, Win—Stay/Lose—Shift (WSLS), Perseveration, Probability Matching and Ideal Predictor (red lines). Each agent performed two repeated binary tasks, one for each Markov order: M0 (left) and M2 (right). The accuracies of the surrogate sequences are also shown (teal lines). Error bars are ± 1 standard deviation.

Regarding the comparison between the output sequences and their surrogates, Figure 3 shows us that the only substantial discrepancy occurs in WSLS and the Ideal Predictor under the M2 task. Again, this is an expected result because those output sequences are the only ones that have underlying structures that are correlated with the input sequences, therefore being susceptible to the action of random shuffling. For the Alternation, Perseveration and Probability Matching behaviours, a random permutation of the output sequence’s elements does not reduce the correlation with the input sequences and therefore does not affect the resulting accuracy or any other first—order statistics.

### 3. Cross—correlation function

The correlation between two sequences temporally shifted from one another by a variable time lag τ is given by the cross—correlation function (CCF). In contrast to the autocorrelation function, the CCF neither yields a unitary coefficient for τ = 0 nor exhibits symmetry around τ = 0. These characteristics are well illustrated in Figure 4: two different, non—correlated sequences, such as an input sequence (either M0 or M2) and an output sequence that bears no causal relation to its respective input (e.g., Alternation, Perseveration or Probability Matching behaviours), will produce a flat CCF (even for the coefficient calculated in τ = 0), therefore being indistinguishable from the CCFs of their surrogates (all panels in Figure 4 except the top—right one). As for the Win—Stay/Lose—Shift behaviour (top—right panel in Figure 4), the CCF is able to reveal its most important trait, namely the great similarity between the input and output sequences for one—step lag. This aspect is captured by the presence of a peak in the CCF for τ = 1, revealing the almost complete correspondence between these sequences when they are one step apart from each other (with the output sequence lagging the input sequence by one time step). The observation of a distinct peak at τ = 1 and an otherwise flat curve in the cross—correlation function between an output sequence and a serially independent input sequence strongly suggests that the output sequence was generated either by a WSLS strategy or, indistinguishably, by simple replication of the input sequence.

**Figure 4.**
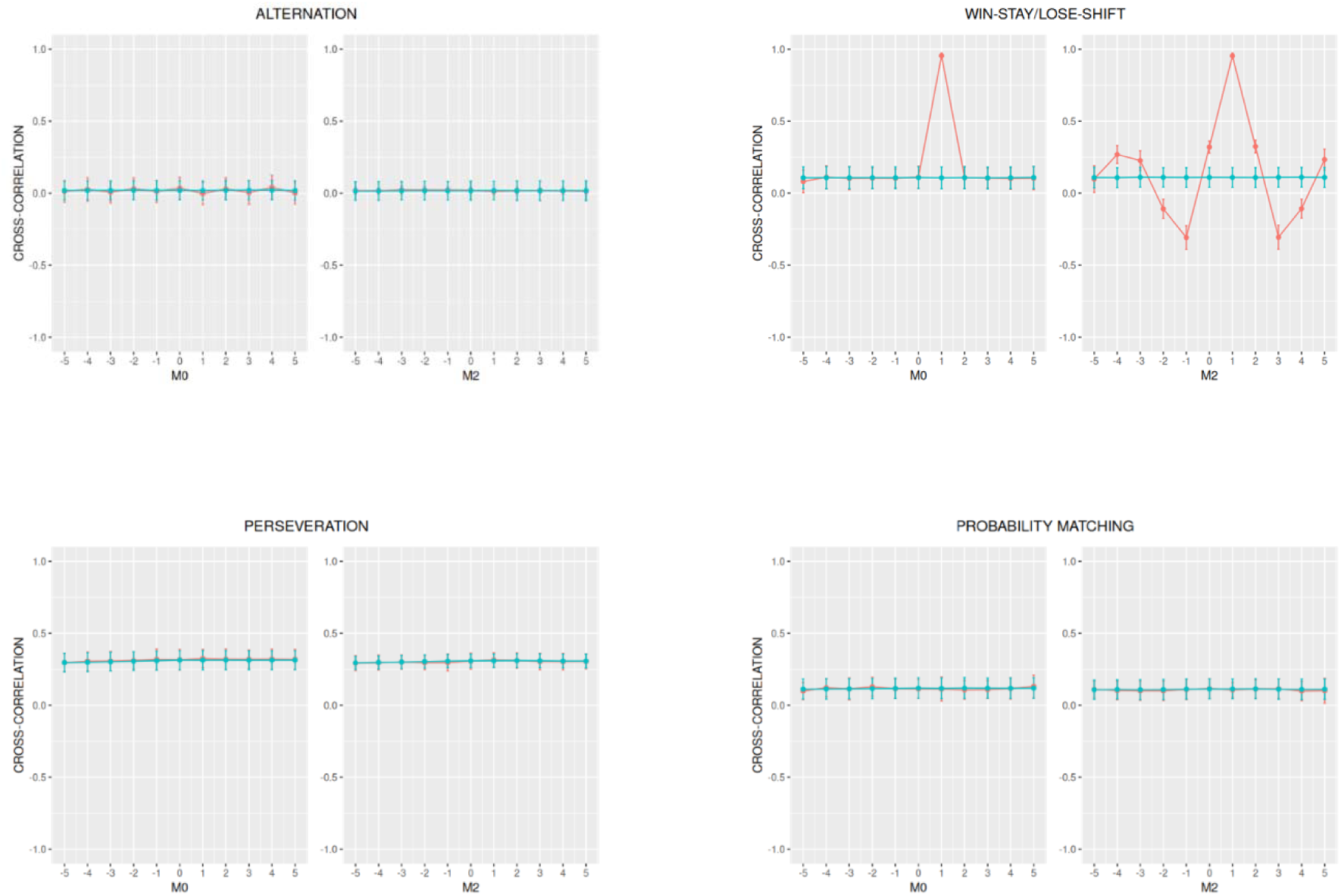
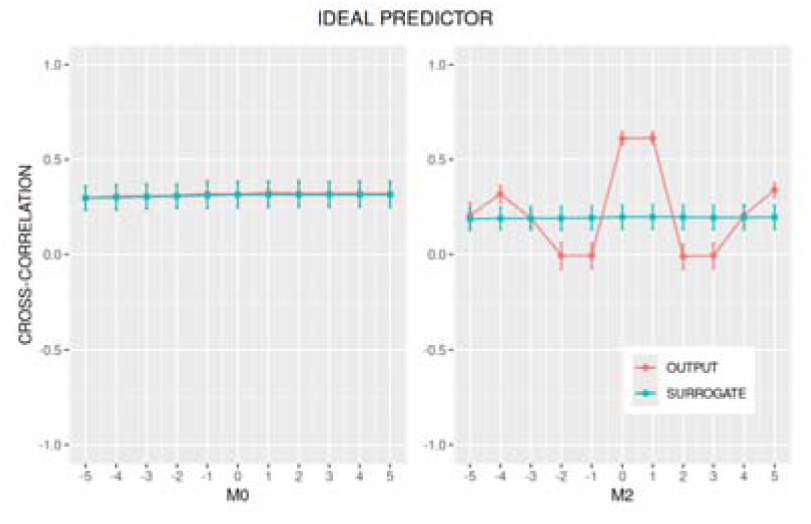
For each behavioural strategy (Alternation, Win—Stay/Lose—Shift (WSLS), Perseveration, Probability Matching, and Ideal Predictor) and for two input Markov sequences (M0 and M2), the mean cross—correlation of the input and the original output sequences is shown as a function of the time lag (red curves), averaged over 50 independent realisations generated under identical conditions. The teal curves represent a grand—mean cross—correlation function derived from surrogate data. For each realisation of the output sequence, 100 surrogate sequences were generated by random permutation. The cross—correlation functions were obtained by first averaging across the 100 surrogates and then averaging across the 50 realisations. In some panels, the cross—correlation curve of the original output sequence is not visible because it is fully superimposed on the corresponding surrogate curve. Error bars indicate ±1 standard deviation.

When the input sequence is generated by a second—order Markov matrix, the interpretation of its cross—correlation with an output sequence that simply mirrors the input sequence — resulting from either a replicating or a WSLS strategy — may be more involved. A distinct peak at τ = 1 will still be evident; however, because the output sequence reproduces a more temporally structured series, the cross—correlation may exhibit peaks at other lag times and an oscillatory pattern, the discussion of which lies beyond the scope of this work.

When an Ideal Predictor responds to an input sequence generated by a zeroth—order Markov process, its strategy coincides with that of a Perseverator; accordingly, their cross—correlation functions are indistinguishable. In contrast, when the input sequence is generated by a second—order Markov process, the resulting cross—correlation function typically exhibits an oscillatory profile, with prominent peaks at multiple time lags. Even in the idealised scenario in which the output of a prototypical behaviour is generated computationally, the interpretation of cross—correlation functions remains non—trivial. Nevertheless, several important points can be highlighted. First, a complex cross—correlation profile — characterised by oscillatory patterns and/or multiple peaks across time lags — emerges only when the two time series share meaningful temporal structure. Second, in the present context, there are two plausible mechanisms capable of generating such temporal relationships: (i) behaviours that largely reproduce the input sequence, as is the case for both WSLS and pure replication strategies; or (ii) at least partially correct prediction of the input sequence by the responding agent, which necessarily implies some form of learning. Although discriminating between these alternatives may be difficult — or even impossible — when relying exclusively on the cross—correlation analysis, a clearer interpretation can be obtained through the combined use of multiple analytical approaches, as proposed in this work.

The symmetry of the CCF around τ = 0 (or, conversely, its absence) is a valuable feature signalling a possible causal relationship between the input and output sequences, beyond the mere temporal correlation between them. A symmetrical CCF means that the correlations between the two sequences are the same whether the output sequence *lags behind* or *leads* the input sequence in time. In contrast, an asymmetrical CCF indicates that those correlations arise in a specific temporal order. From the top—right panel in Figure 4, we can conclude that the high positive peak at τ = 1 in both M0 and M2 (as well as secondary peaks in M2) is produced only when the output sequence is lagging (is in the past of) the input sequence by a given amount of time. Hence, this asymmetry means that the content of the input sequence bears an influence on the decisional process of the agent (human or not) that generates the output sequence.

### 4. Autocorrelation function

As defined previously, the autocorrelation function (ACF) gives the correlation between a sequence and a shifted realisation of itself as a function of the time lag. For a lag τ = 0, the ACF is equal to the correlation between two identical sequences and thus trivially equal to unity: C_Y,Y_(0) ≡ 1. Also, as we can see in Figure 5, the ACF is always symmetrical around τ = 0, i.e., a positive lag τ = +k yields the same value as τ = —k. From Figure 5 we can also appreciate that the ACF is able to reveal relevant features of the underlying sequence. In the Alternation behaviour (Figure 5, top—left panel), the function heavily oscillates, reflecting the high frequency of alternating choices, and fades away with increasing lags (in absolute value). This reduction in the ACF’s envelope is due to the small amount of noise (5%) introduced into the agent’s probability of alternating between choices. In that same panel, we can notice how the shuffling of the output sequences destroys any serial correlation previously present in the sequences, bringing all autocorrelation coefficients to zero (except, of course, for τ = 0). As a matter of fact, the output sequences generated by the Alternation behaviour are well described by a first—order Markov chain, a feature that is captured by the profile of their ACF, in which the oscillating period is 2.

**Figure 5.**
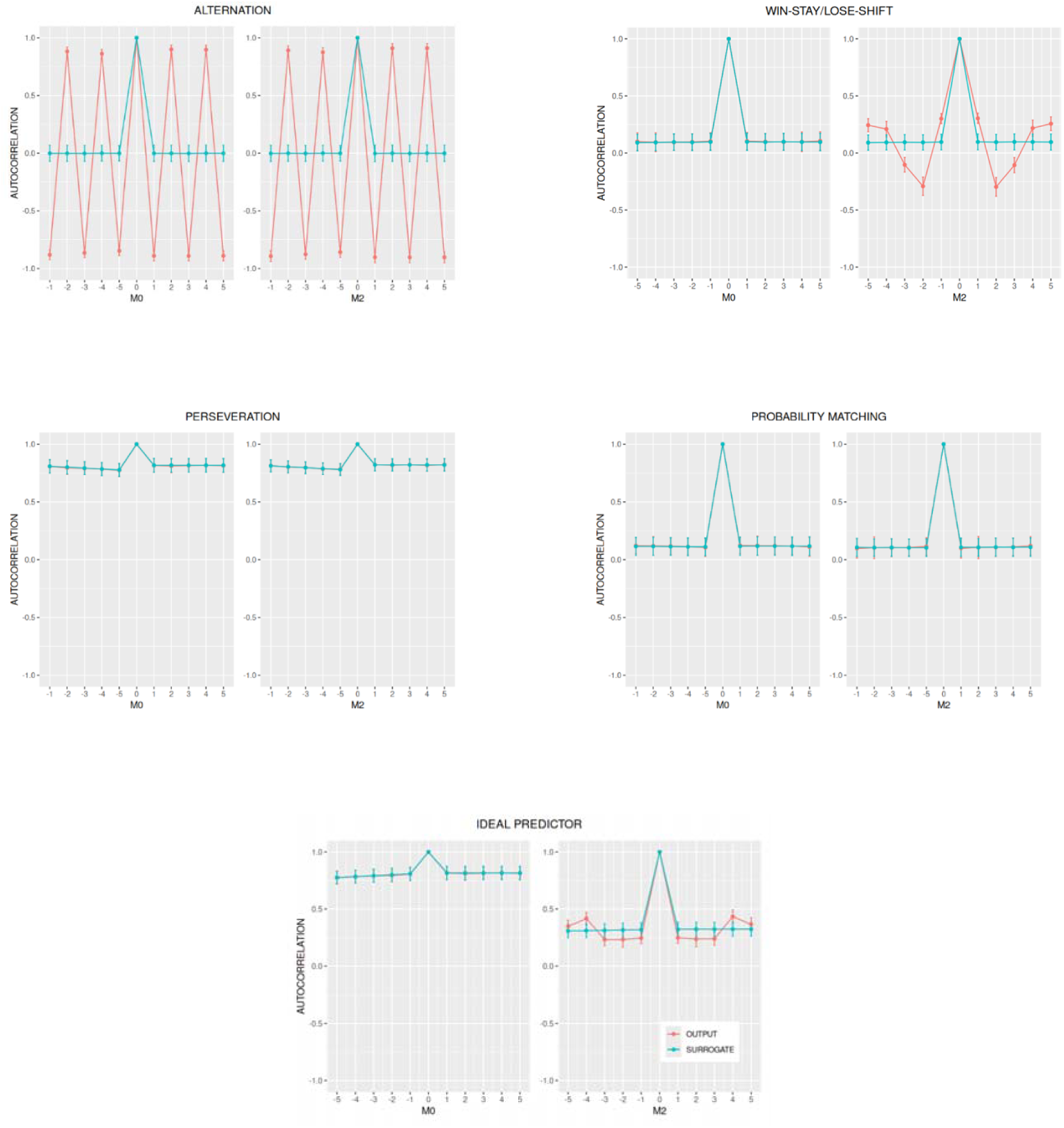
For each behavioural strategy (Alternation, Win—Stay/Lose—Shift (WSLS), Perseveration, Probability Matching, and Ideal Predictor) and for two input Markov sequences (M0 and M2), the mean autocorrelation of the original output sequences is shown as a function of the time lag, averaged over 50 independent realisations generated under identical conditions. The teal curves represent a grand—mean autocorrelation function derived from surrogate data. For each realisation of the output sequence, 100 surrogate sequences were generated by random permutation. The autocorrelation functions were obtained by first averaging across the 100 surrogates and then averaging across the 50 realisations. In some panels, the autocorrelation curve of the original output sequence is not visible because it coincides with the corresponding surrogate curve. Error bars indicate ±1 standard deviation.

In the Probability Matching behaviour (Figure 5, bottom—right panel), the intrinsically random output sequence is associated with a very low serial correlation, which is manifested by a flat ACF. The above—zero baseline is due to the unequal proportions of +1s and −1s in the sequence (0.67 and 0.33, respectively) as well as the non—normalised definition of the ACF we adopted in the present work (Figure 5 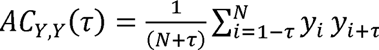, *for τ* = 0, −1, −2, … , −(*N* − 1) (Equation 12Equations 11 and 12). The same flat ACF is seen in Perseveration (Figure 5, bottom—left panel), although with a much higher baseline. The reason for this is that each output sequence is composed almost exclusively of +1s with a small amount (5%) of −1s randomly scattered throughout. Finally, the top—right panel in Figure 5 shows the ACF for the Win—Stay/Lose—Shift behaviour. Since this behaviour essentially replicates the input sequence, the ACF for the output sequence is essentially the autocorrelation of the input sequence itself. As a result, the ACF in M0 returns a flat curve, whereas in M2, the ACF reflects the higher content of serial correlation present in a second—order Markov sequence. Again, in the surrogate sequences, any serial correlation is removed by the random shuffling.

### 5. Shifting probabilities

These measures evaluate the probability of an agent shifting their current choice in relation to the previous outcome. Whereas *P*(shift) provides a measure of the unconditional probability of shifting, *P*(shift|win) and *P*(shift|lose) provide a measure of the conditional probability of shifting given that the previous choice led to a win (success) or to a loss (failure), respectively. For the prototypical behaviours under scrutiny here, we observe: (i) a high probability of shifting in Alternation, in both unconditional and conditional measures, since it represents the ultimate example of exploratory/shifting behaviour; (ii) a high *P*(shift|lose) and low *P*(shift|win) in WSLS, which, by construction, is a behaviour characterised by the agent staying (not shifting) when it wins and shifting when it loses; (iii) a low probability of shifting in Perseveration, for both unconditional and conditional measures, since it represents, in opposition to Alternation, the most extreme variety of exploitative/staying behaviour; (iv) no large deviations from 0.5 for all shifting probability measures in the Probability Matching behaviour, since it essentially generates random sequences that bear no correlation between their shifts and the “win” or “lose” Accuracy). Therefore, we observe, M0 or M2, similar to either the Perseveration’s or the WSLS’s results, respectively.

The process of generating surrogate sequences, by removing serial correlations from the original output sequences, impacted the shifting probabilities mainly for Alternation, Win—Stay/Lose—Shift, and Ideal Predictor for the M2 condition, whose sequences carry a serial structure. By the same token, the random shuffling that creates the surrogate sequences spared Perseveration, Probability Matching and Ideal Predictor (in the M0 condition) from noteworthy changes in their shifting probabilities, since there is no serial correlation in those agents’ responses. Indeed, all of these observations are evident in Figure 6 and 7, which show the shifting probabilities for the original output and surrogate sequences, for all five behaviours, under both Markov conditions.

**Figure 6.**
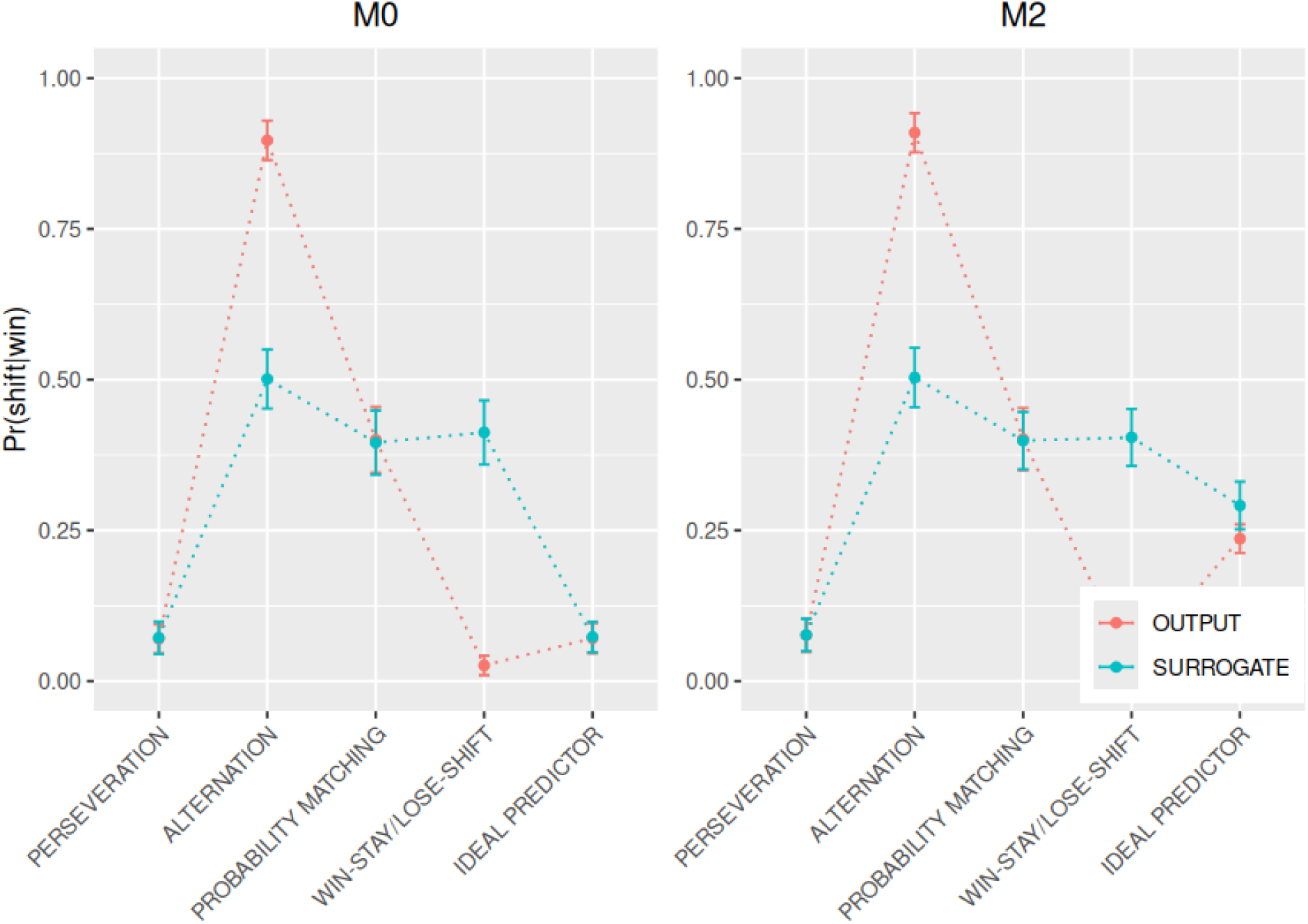
Average shifting probabilities, calculated for the original output sequences (in red) and their respective surrogates (in teal) for both conditions (M0 and M2). Each curve is an average over 50 output sequences for each behaviour (Alternation, Win—Stay/Lose—Shift (WSLS), Perseveration, Probability Matching and Ideal Predictor). A set of 100 surrogate sequences was obtained from each original output sequence by a random permutation of its elements. After calculating the mean shifting probability for each set of 100 output surrogates, a grand mean was obtained by averaging them over the set of 50 mean values, separately for each order and behaviour (teal lines). The figure shows, the conditional probability of shifting given that the agent’s previous choice led to a “win”, P(shift|win). Error bars are ± 1 standard deviation.

**Figure 7.**
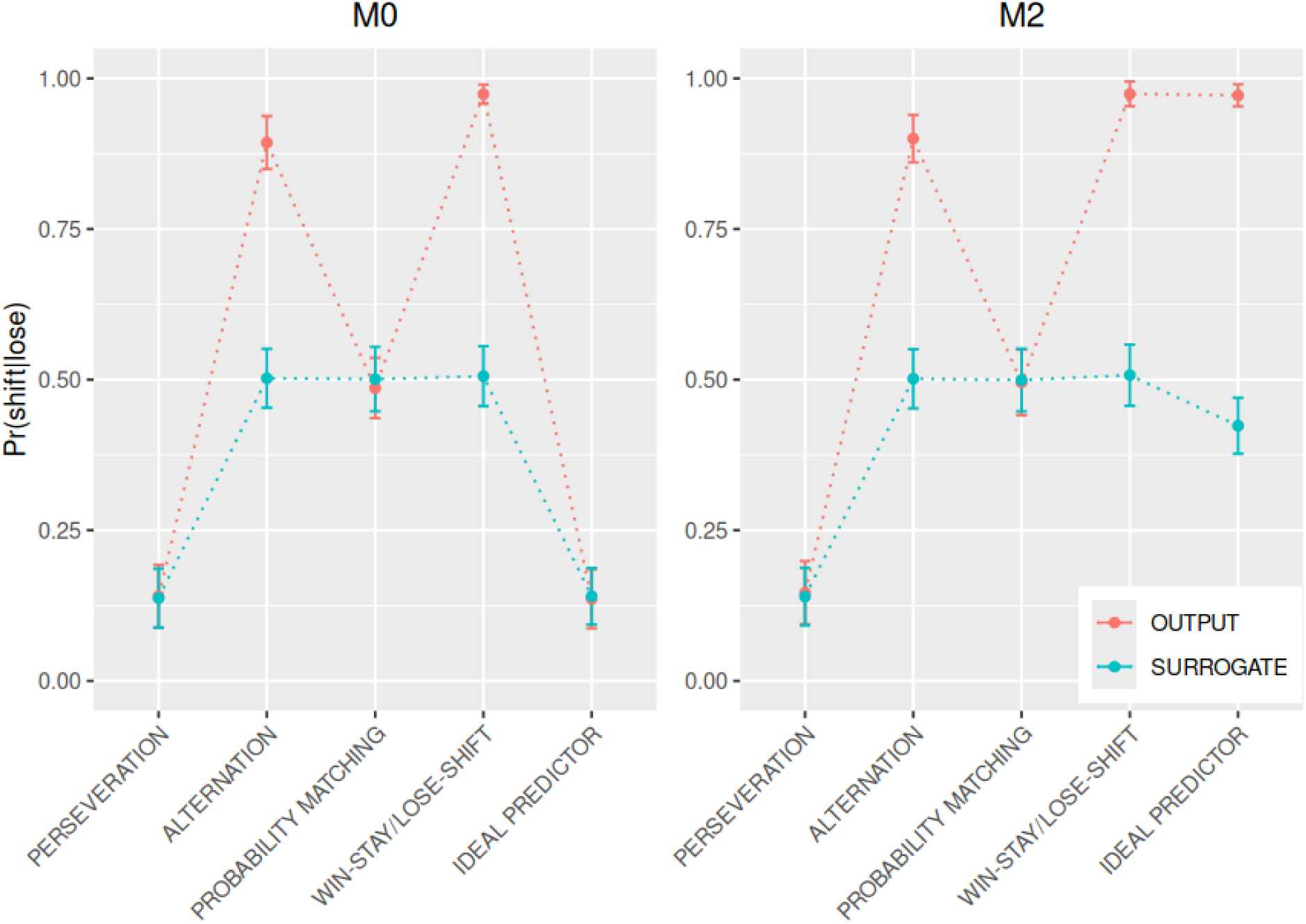
Average shifting probabilities, calculated for the original output sequences (in red) and their respective surrogates (in teal) for both conditions (M0 and M2). Each curve is an average over 50 output sequences for each behaviour (Alternation, Win—Stay/Lose—Shift (WSLS), Perseveration, Probability Matching and Ideal Predictor). A set of 100 surrogate sequences was obtained from each original output sequence by a random permutation of its elements. After calculating the mean shifting probability for each set of 100 output surrogates, a grand mean was obtained by averaging them over the set of 50 mean values, separately for each order and behaviour (teal lines). The figure shows the conditional probability of shifting given that the agent’s previous choice led to a “lose”, P(shift|lose). Error bars are ± 1 standard deviation.

Plotting *P*(shift|win) versus *P*(shift|lose) in the shifting diagram for the zeroth—order Markov chain (Figure 8) allows behaviours to be distinguished by quadrant. Four of the strategies (Alternation, Ideal Predictor, Perseveration and Win—Stay/Lose—Shift) are located at the corners of the quadrants, indicating the type of strategy associated with each behaviour. Only Probability Matching is in the centre of the diagram. An almost identical pattern appears in Figure 9, the shifting diagram for the second—order Markov chain, except that the Ideal Predictor has migrated to the Win—Stay/Lose—Shift quadrant. This is because, as remarked earlier, the specific M2 sequences generated for this study favour a Win—Stay/Lose—Shift behaviour.

**Figure 8.**
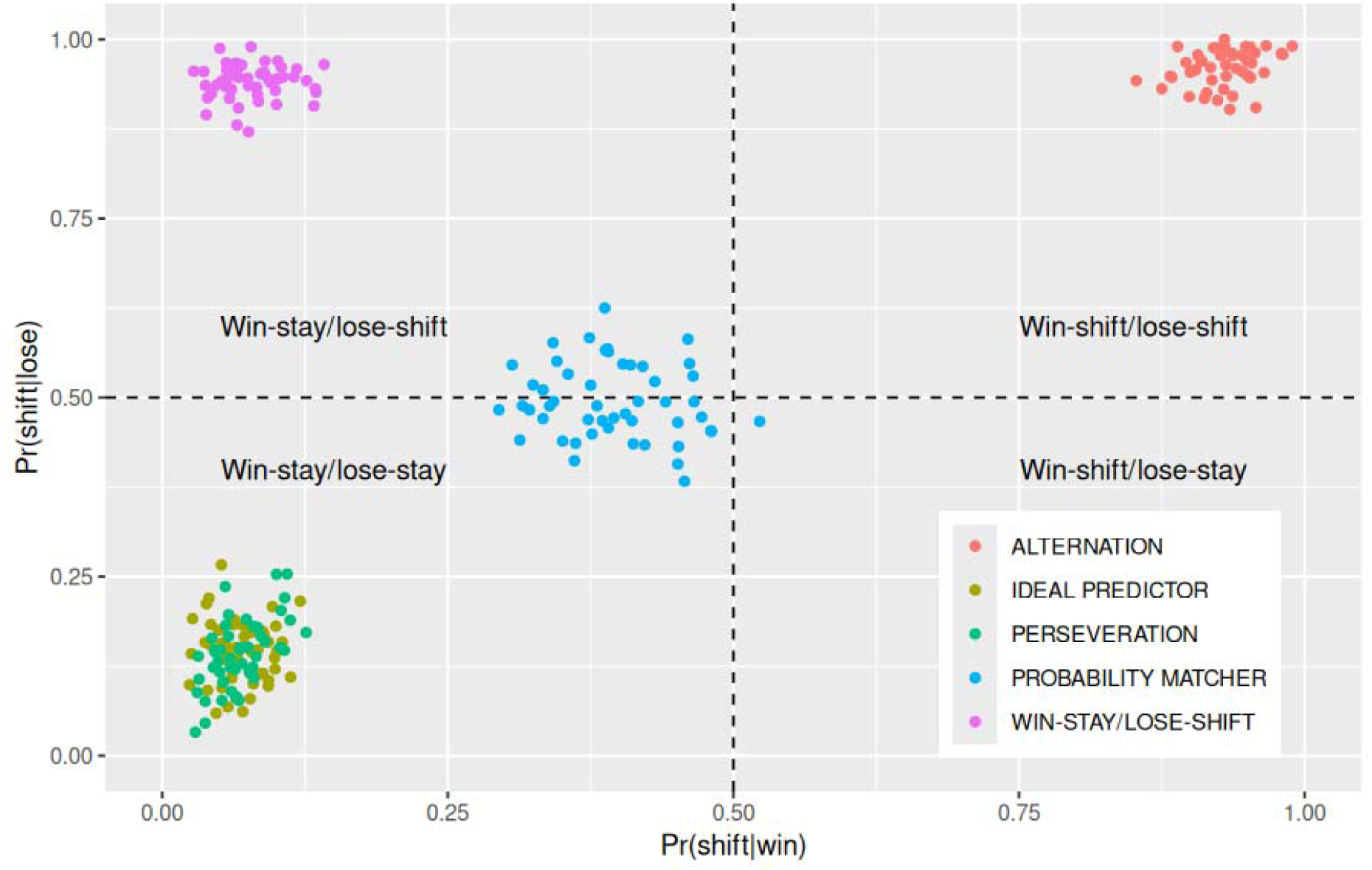
Scatterplot showing the probability of shifting given an incorrect prediction, P(shift|win) versus the probability of shifting given a correct prediction, P(shift|lose) for the M0 condition.

**Figure 9.**
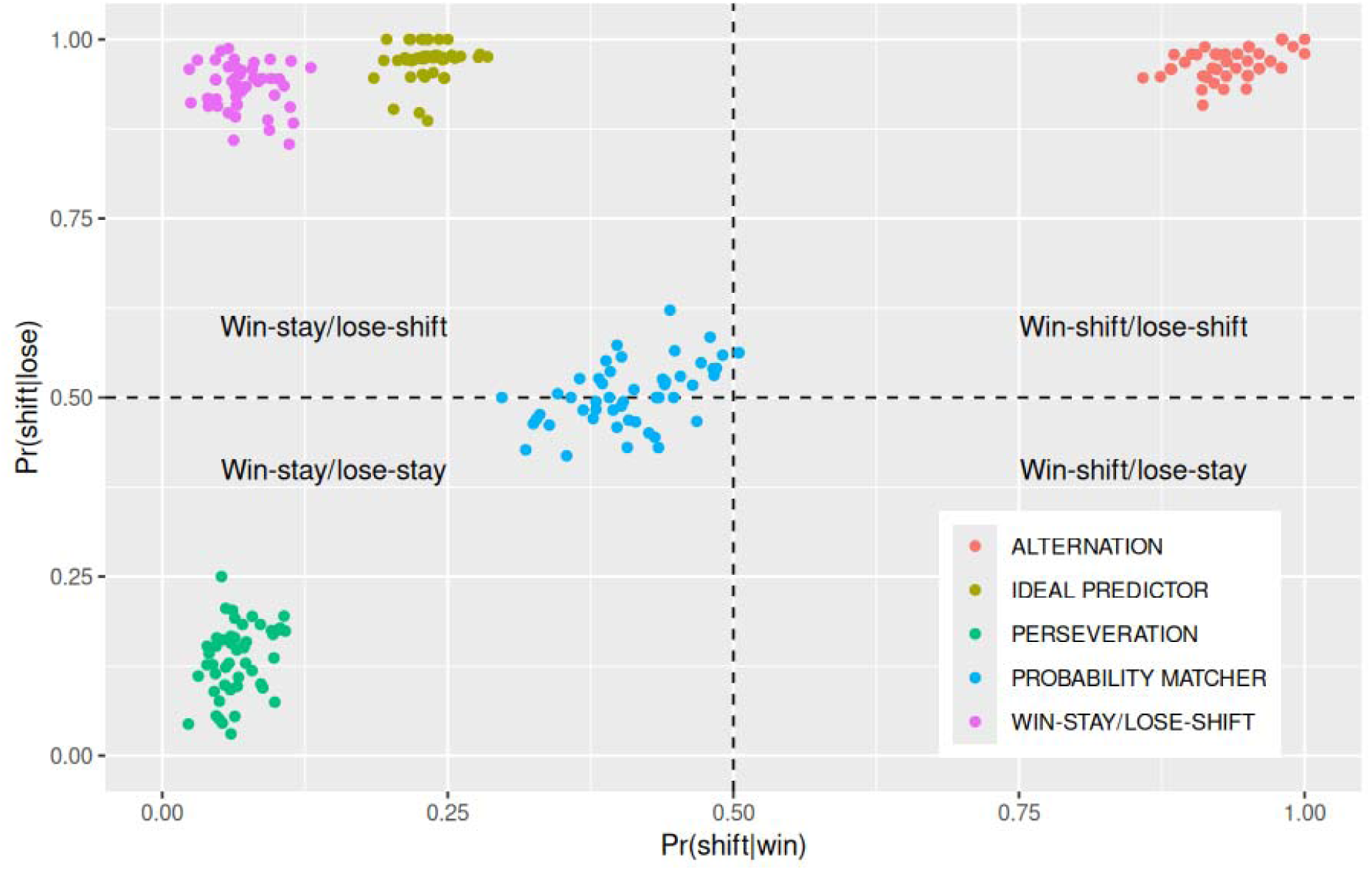
Scatterplot showing the probability of shifting given an incorrect prediction, P(shift|win) versus the probability of shifting given a correct prediction, P(shift|lose) for the M2 condition.

### 6. Reconstruction of Markov matrices

For this analysis, as shown in Table IV, we first computed the matrices that reflect, for each behaviour in the M0 condition, the asymptotic probabilities of each binary event in their respective output sequences, consistent with the mean responses, as depicted in Figure 2 (Simulation of Prototypical Behaviours — Results and Partial Discussion).

**Table IV.** Zeroth—order Markov matrices reconstructed from the output sequences generated by the five simulated behaviours (Alternation, Win—Stay/Lose—Shift, Perseveration, Probability Matching and Ideal Predictor) in response to an M0 input sequence. The value of each element in the matrices is expressed by a ± 1 standard deviation calculated from the set of 50 virtual agents simulated in each behaviour.

| ALTERNATION |  | WIN—STAY/LOSE—SHIFT |  |
| --- | --- | --- | --- |
| | $\begin{array}{c\vert cc} & \begin{matrix} y_i \\ -1 \quad +1 \end{matrix} \\ \hline \begin{matrix} x_{i-1} \\ \hline \end{matrix} & [0.49, 0.51] & [0.49, 0.51] \end{array}$ | | $\begin{array}{c\vert cc} & \begin{matrix} y_i \\ -1 \quad +1 \end{matrix} \\ \hline \begin{matrix} x_{i-1} \\ \hline \end{matrix} & [0.31, 0.38] & [0.62, 0.69] \end{array}$ |
| PERSEVERATION |  | PROBABILITY MATCHING |  |
| | $\begin{array}{c\vert cc} & \begin{matrix} y_i \\ -1 \quad +1 \end{matrix} \\ \hline \begin{matrix} x_{i-1} \\ \hline \end{matrix} & [0.04, 0.07] & [0.93, 0.96] \end{array}$ | | $\begin{array}{c\vert cc} & \begin{matrix} y_i \\ -1 \quad +1 \end{matrix} \\ \hline \begin{matrix} x_{i-1} \\ \hline \end{matrix} & [0.29, 0.37] & [0.63, 0.71] \end{array}$ |
| IDEAL PREDICTOR |  |  |  |
| | $\begin{array}{c\vert cc} & \begin{matrix} y_i \\ -1 \quad +1 \end{matrix} \\ \hline \begin{matrix} x_{i-1} \\ \hline \end{matrix} & [0.04, 0.06] & [0.94, 0.96] \end{array}$ | | |

We also computed the matrices in Table V, which reflect, for each behaviour in the M2 condition, the probabilities of each binary event in the output sequences conditionally on the history of the input sequence. The matrices in Table V, although more involved than those in Table IV, are of easy interpretation. In Alternation (top—left matrix), the confidence intervals for all elements in the matrix include the value 0.50, which, in this behaviour, is the probability of occurrence of both +1 and −1 in the output sequence, independently of the preceding events in the input sequence. In Win—Stay/Lose—Shift, as shown in Table V (top—right matrix), whenever an event (either +1 or −1) in the output sequence (y_i_) is immediately preceded by the same event in the input sequence (x_i—1_), the confidence interval for that matrix element always encloses the value 0.95 (the probability of staying given a win or shifting given a loss), irrespective of the second previous event (x_i—2_). This result reflects the identity between the Win—Stay/Lose—Shift behaviour and mere replication of the input sequence.

**Table V.**
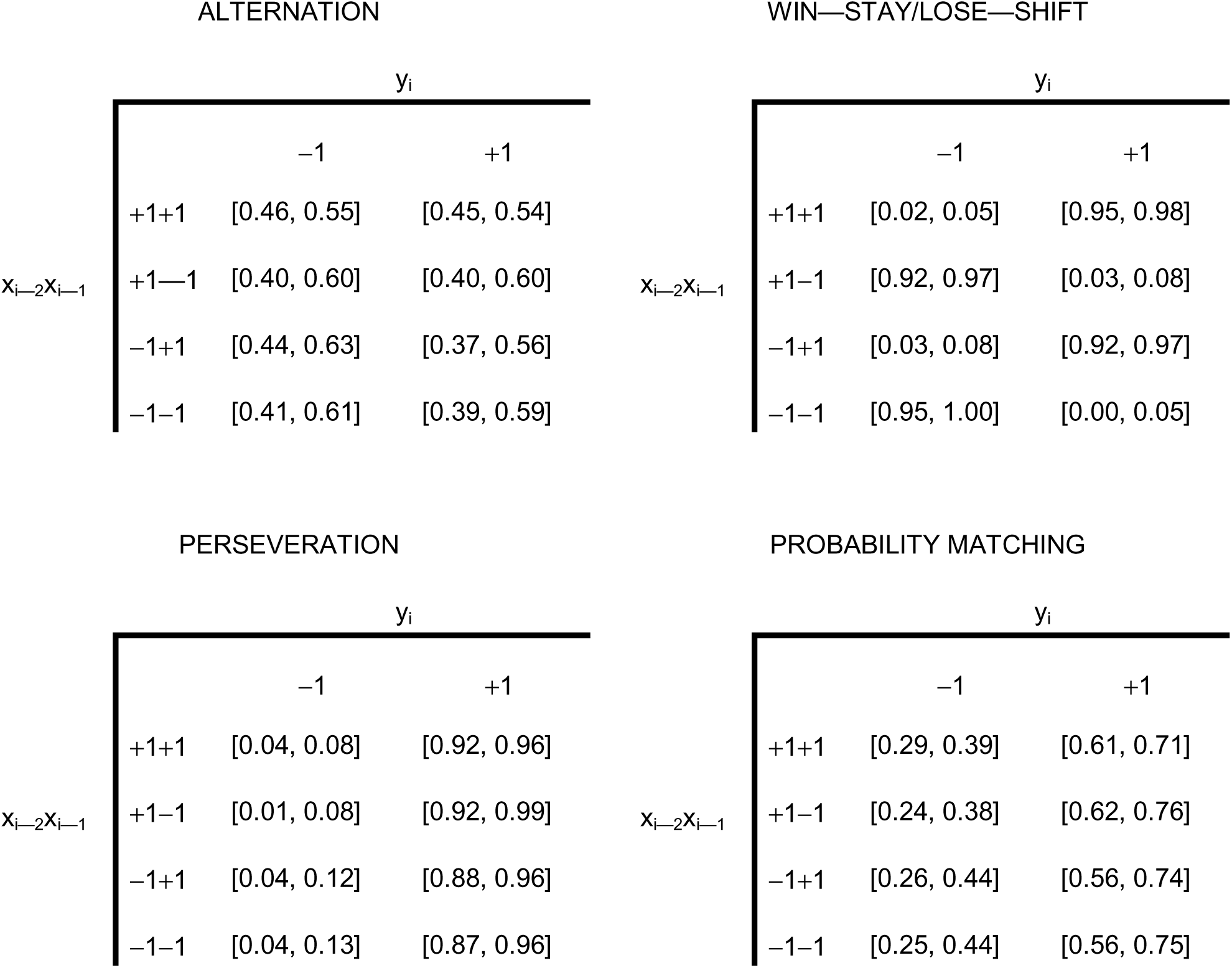
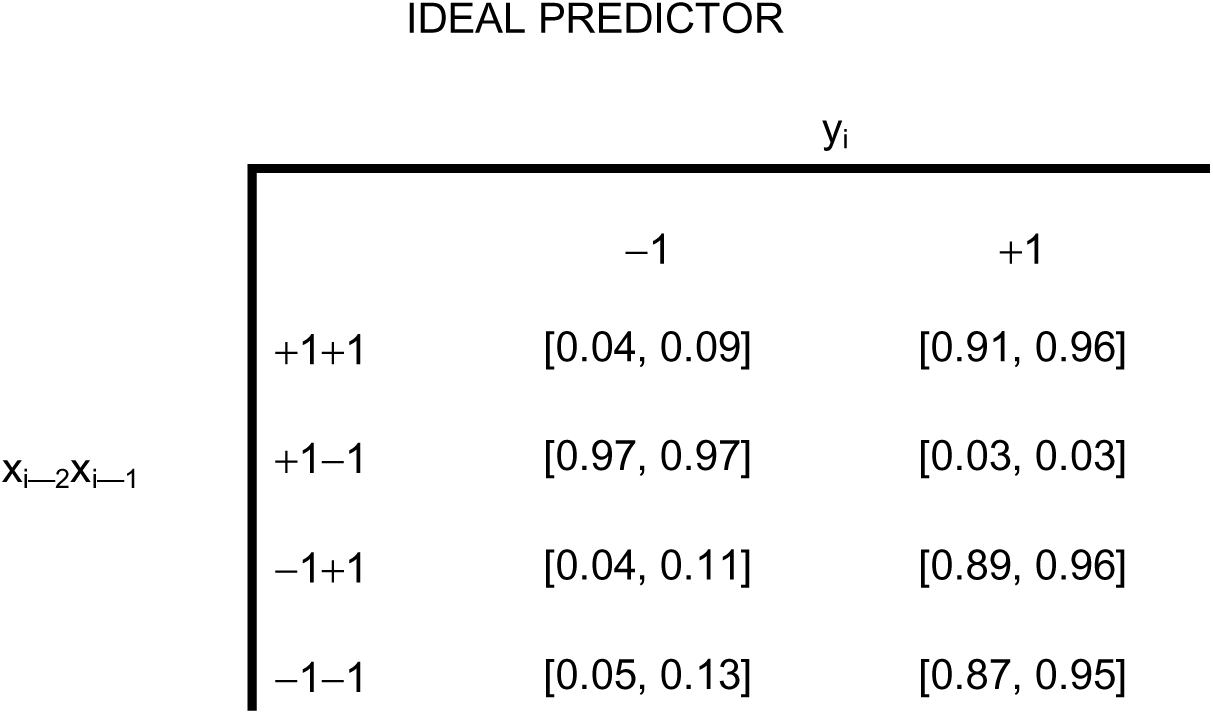
Second—order Markov matrices reconstructed from the output sequences generated by the five simulated behaviours (Alternation, Win—Stay/Lose—Shift, Perseveration, Probability Matching and Ideal Predictor) in response to an M2 input sequence. The value of each element in the matrices is expressed by a ± 1 standard deviation calculated from the set of 50 virtual agents simulated in each behaviour.

The confidence intervals in Table V also show that, for the Perseveration behaviour, irrespective of the previous elements in the input sequence, all entries in the matrix reflect a concentration of choices on the most likely result (y_i_ = +1), whereas for the Probability Matching behaviour, they exhibit a distribution of probabilities across the two columns that mirrors the first—order statistics of the input sequence. As for the Ideal Predictor, Table V shows that a “perseveration” strategy is also present, but it is no longer grounded in the simpler first—order statistics of the input sequence; rather, it relies on the conditional probabilities of an outcome given the two preceding input events x_i—1_ and x_i—2_, as explained in Description of prototypical behaviours.

Even though the reconstructed Markov matrices may share a great deal of information with other measures, such as the mean response and shifting probabilities, they offer an alternative and useful perspective to interpret the decisional strategy adopted by the agent in a binary choice task. An important distinctive feature of the Markov reconstruction is the information it can provide concerning the “memory” of the sequence, an aspect that is not addressed by the other measures.

The results in Table V are also shown in Figure 10. This graph reflects how each behaviour responds to the sequence to be predicted, depending on the elements of the Markov matrix of the input sequence. Qualitatively speaking, it can compare the distance between the mean prediction for each agent type (for example, the Ideal Predictor chooses +1 after +1+1) and the respective element of the input transition matrix. This distance, and the distance for the other three elements, can give us an idea of how an agent responded to the input sequence.

**Figure 10.**
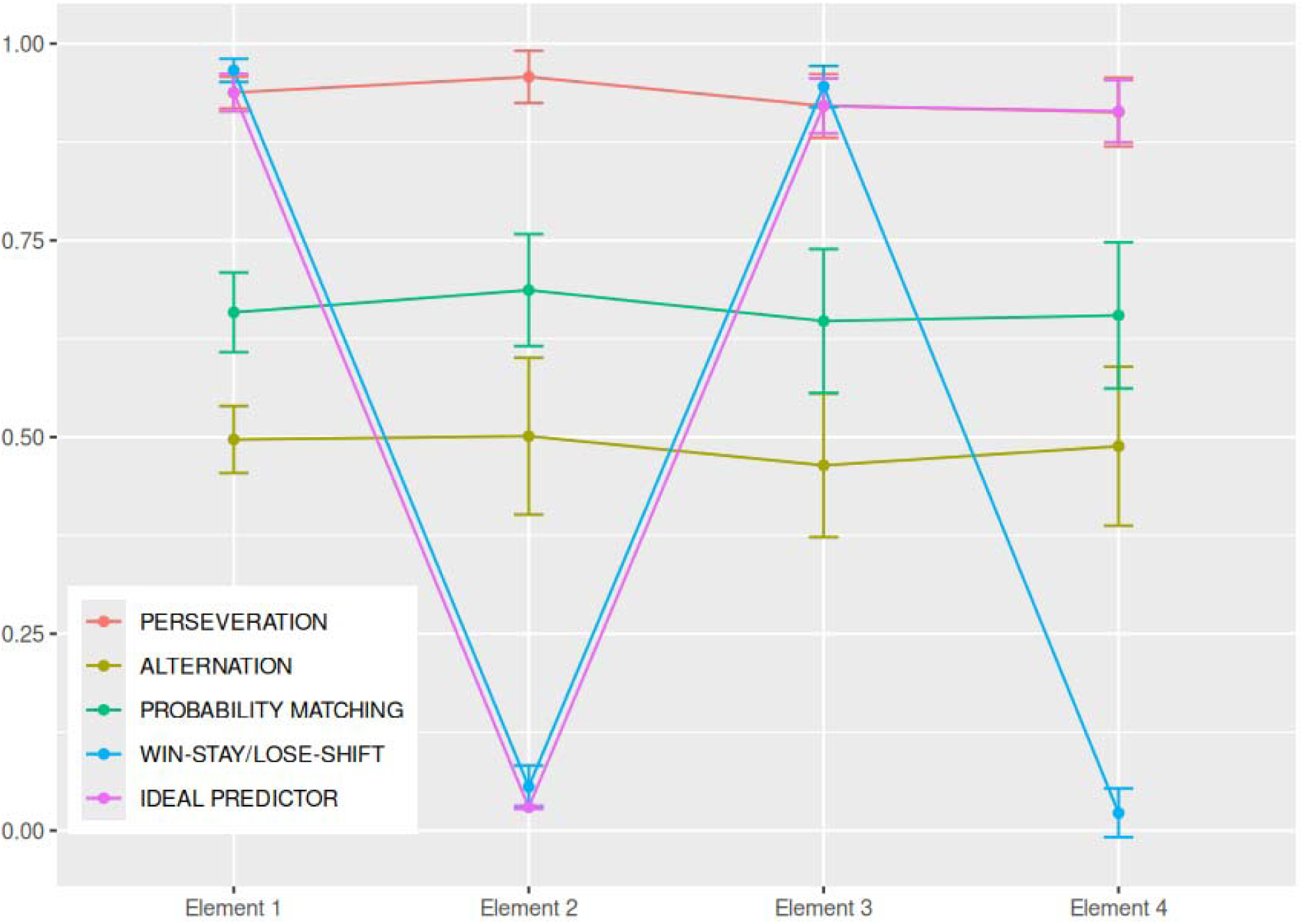
Graph showing the four elements of the Markov reconstructed matrix for each of the five behaviours (Alternation, Win—Stay/Lose—Shift (WSLS), Perseveration, Probability Matching and Ideal Predictor) under the second—order Markov sequence. Error bars are ± 1 standard deviation.

## Application of the Analytical Tools to Empirical Data

Here we will present the use of Markov—generated input sequences and the analytical tools previously described to study decisional behaviours of human participants during repeated binary choice tasks in a visual probability learning setting. Our aim was to apply the analytical tool kit to a simple procedure where the experimental manipulation would be already known to produce sizable effects on the dependent variables. To this end, the elected procedure involved the differential effect of gains and losses as payoffs in a binary choice task (Bereby—Meyer & Erev, 1998; Erev et al., 1999; Myers et al., 1961, 1963; Myers & Suydam, 1964). The main idea was to evaluate the efficacy of the present techniques in extracting from the binary sequences as much of the available information as possible, as well as any structural element that could reveal a differential effect of the payoffs.

### 1. General experimental procedure

In both experiments, the participant sat comfortably in front of a 24—inch computer monitor with a screen resolution of 1920 × 1080 at 60 Hz, and the keyboard was used by the volunteer to enter their prediction on each trial. In each session, a trial began with the presentation of a screen containing the general instructions, after which a screen showing the request “Choose either right or left” remained until one of the horizontal arrow keys on the keyboard had been pressed by the participant. Following the key press, the image of a five pence coin, in Experiment I, or an “X” in Experiment II, was presented either on the left or on the right side of the screen, corresponding to the current input sequence event, along with the message “Congratulations!” or “Sorry!…”, depending on the participant’s prediction being correct or incorrect (see Figure 11 and Figure 12). In both experiments, the participants were randomly assigned to two groups (Win and Lose), and no blinding procedures were implemented during either data collection or data analysis. Inclusion criteria: (i) volunteers of either sex; (ii) aged between 20 and 55 years; (iii) good physical and mental health; (iv) genuine willingness to participate in the study.

**Figure 11.**
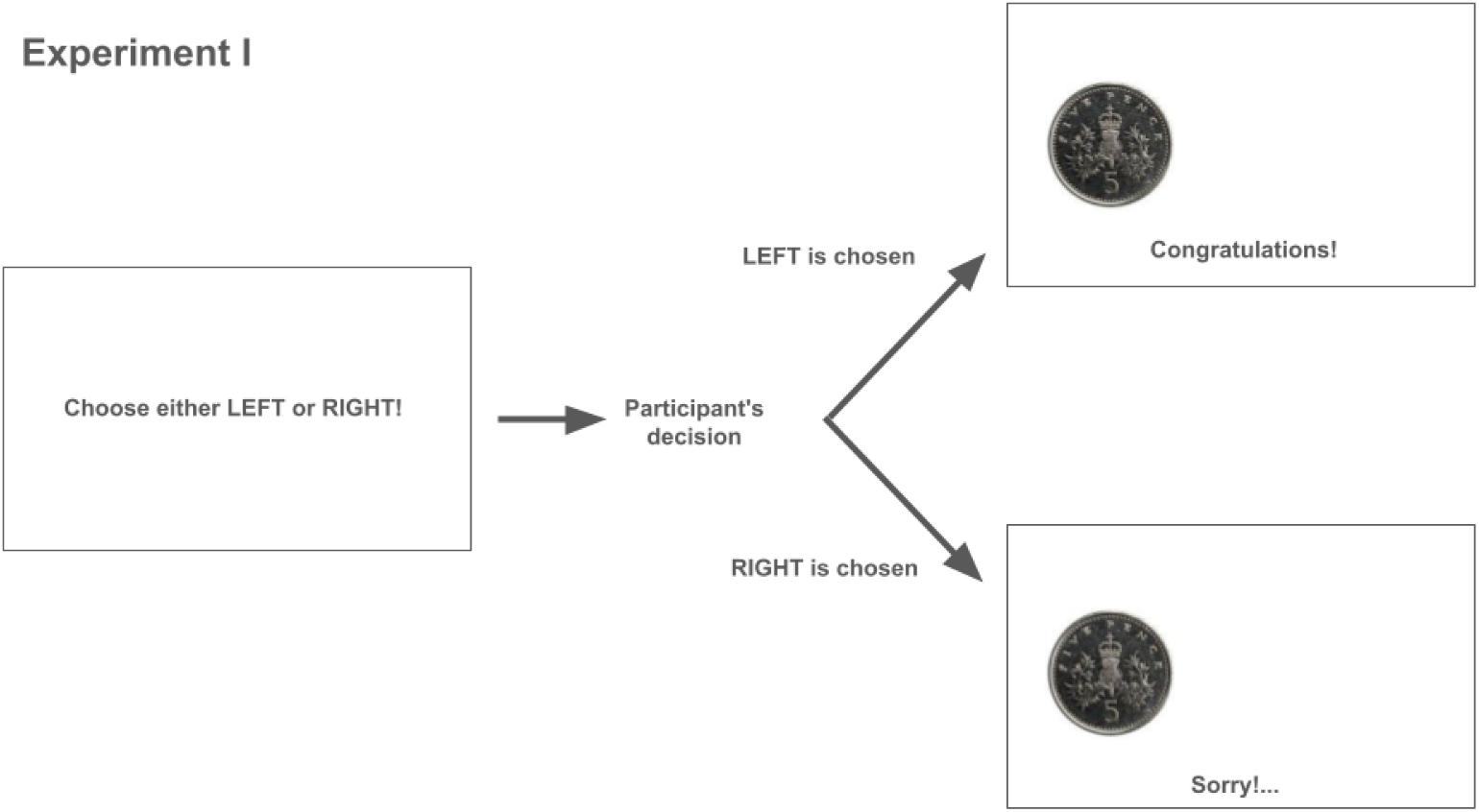
Schematic view of the task sequence in Experiment I. In each trial a screen prompted the participant to predict one out of two designated locations (right or left) for the presentation of the visual stimulus (the image of a five pence coin), by pressing one of the horizontal arrow keys on the keyboard; after the participant’s choice, the result provided by the input sequence was shown on the next screen (in this example, the binary result in that trial was “left”); if the participant’s prediction was correct (a “win”), the feedback message “Congratulations” was displayed, otherwise the message was “Sorry!…”.

**Figure 12.**
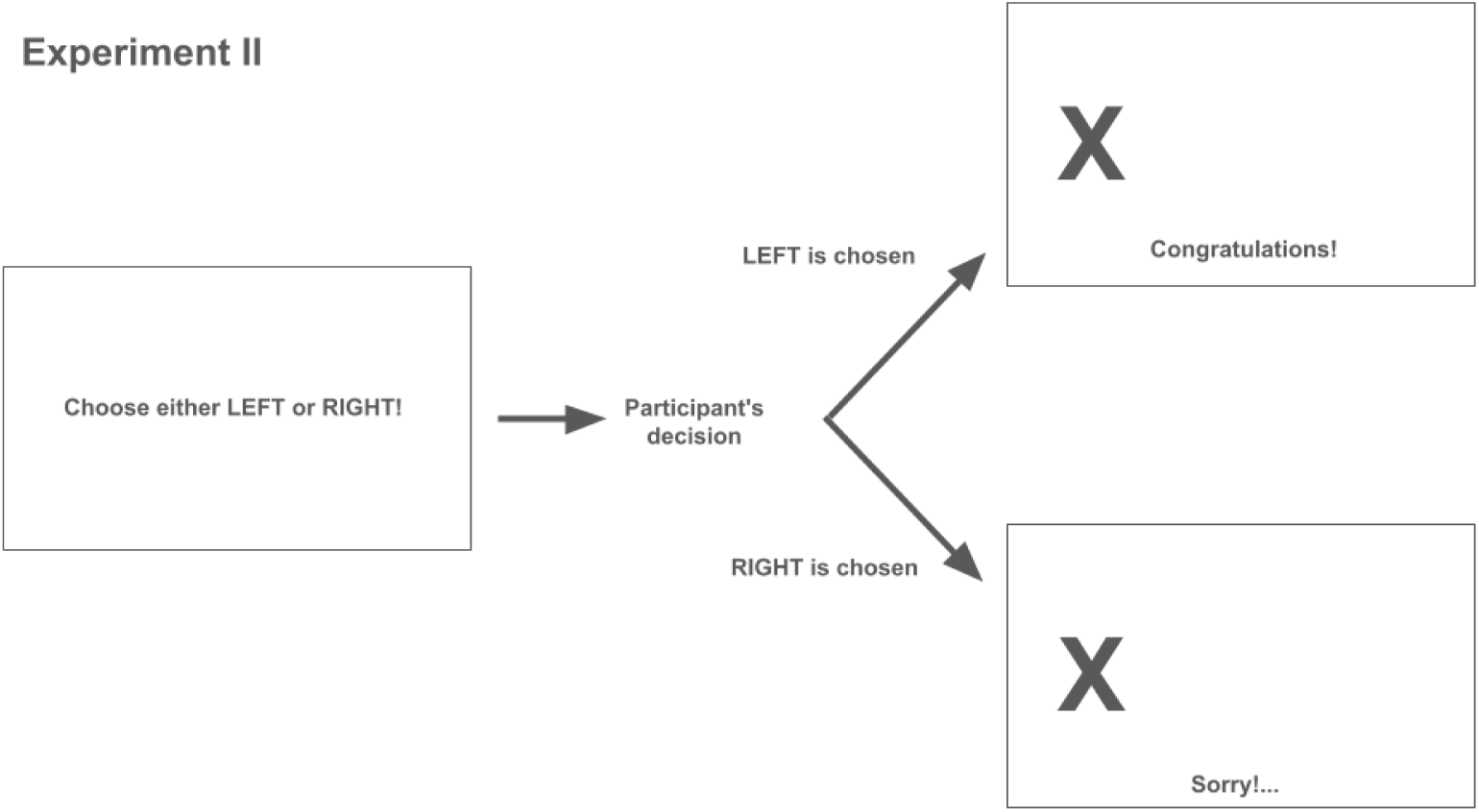
Schematic view of the task sequence in Experiment II. In each trial a screen prompted the participant to predict one out of two designated locations (right or left) for the presentation of the visual stimulus (the image of an “X”), by pressing one of the horizontal arrow keys on the keyboard; after the participant’s choice, the result provided by the input sequence was shown on the next screen (in this example, the binary result in that trial was “left”); if the participant’s prediction was correct (a “win”), the feedback message “Congratulations” was displayed, otherwise the message was “Sorry!…”.

Exclusion criteria: (i) self—reported neurological and/or cognitive impairments, or use of centrally acting drugs; (ii) voluntary decision to withdraw from the experiment at any time; (iii) inability to complete the experimental procedure in its entirety. The illustrative inferential analyses were conducted after verifying, where necessary, that the relevant statistical assumptions were satisfied.

### 2. Specific procedures

#### a. Experiment I

Twelve volunteers participated in this study (7 females and 5 males, mean age ± s.d.: 25.6 ± 3.9 years). The experiment had a total of 150 trials divided into three successive blocks of equal length; a single input sequence (M0), previously generated with the proportion of +1s equal to 67% and of —1s equal to 33%, was shuffled and used in testing all twelve participants. The experiment was carried out in the Department of Experimental Psychology at the University of Oxford. The experimental procedure was approved by the University of Oxford’s Ethics Committee.

The volunteers were randomly split into two groups of six participants: the participants of the “Win” group were rewarded (by earning 5 pence) for each correct prediction and not penalised for any incorrect one; in the “Lose” group, participants were punished (by losing 5 pence from a pre—determined initial amount) for each incorrect prediction and not rewarded for any correct ones.

#### b. Experiment II

Twenty—eight volunteers participated in this study (13 females and 15 males, mean age ± s.d.: 30.9 ± 8.6 years). The experiment had a total of 100 trials; each volunteer had to predict a different input sequence generated by a second—order Markov chain (see M2, Table III). The experimental procedure was approved by the University of São Paulo’s Ethics Committee.

The volunteers were randomly split into two groups of 14 participants each: the participants of the “Win” group received fifty cents of Brazilian Real (R$) for each correct prediction and were not penalised for any incorrect ones; in the “Lose” group, participants were penalised by fifty cents of Brazilian Real from a pre—determined initial amount of R$5 for each incorrect prediction and not rewarded for their correct ones.

### 3. Results and partial discussion

Although inferential statistical analyses are not required to establish the applicability or utility of the proposed toolkit, they were conducted not as confirmatory hypothesis tests, but rather to illustrate that conventional metrics such as accuracy and mean response may fail to capture more subtle behavioural differences — and the ubiquitous “statistically significant” interpretations that usually follow them — especially under the relatively low statistical power of the hypothesis—testing settings in Experiments I and II. In contrast, under the same conditions, other measures within the toolkit were able to reveal some “statistically significant” effects, as will be shown below (for a critical discussion of “statistical significance”, see Wasserstein et al., 2019. Therefore, the words “significance”, “significant” and “significantly” will be used hereafter in quotation marks to reflect our endorsement of that critical stance).

#### a. Mean response and Accuracy

Figure 13 shows the mean response and accuracy for the two experimental groups, Win and Lose, in both experiments I and II. A Welch’s *t*—test was conducted for each experiment, using each measure (mean response and accuracy) as dependent variables and group (win or lose) as the independent variable. No statistically significant differences between the groups were observed in either experiment (Exp. I, mean response: *t*(9.334) = 0.910, *p* = 0.386, *Cohen’s d* = 0.484; Exp. I, accuracy: *t*(7.152) = 0.258, *p* = 0.386*, Cohen’s d* = 0.803; Exp. II, mean response: *t*(25.357) = 0.400, *p* = 0.630, *Cohen’s d* = 0.138; Exp. II, accuracy: *t*(21.389) = 0.420, *p* = 0.679, *Cohen’s d* = 0.154).

**Figure 13.**
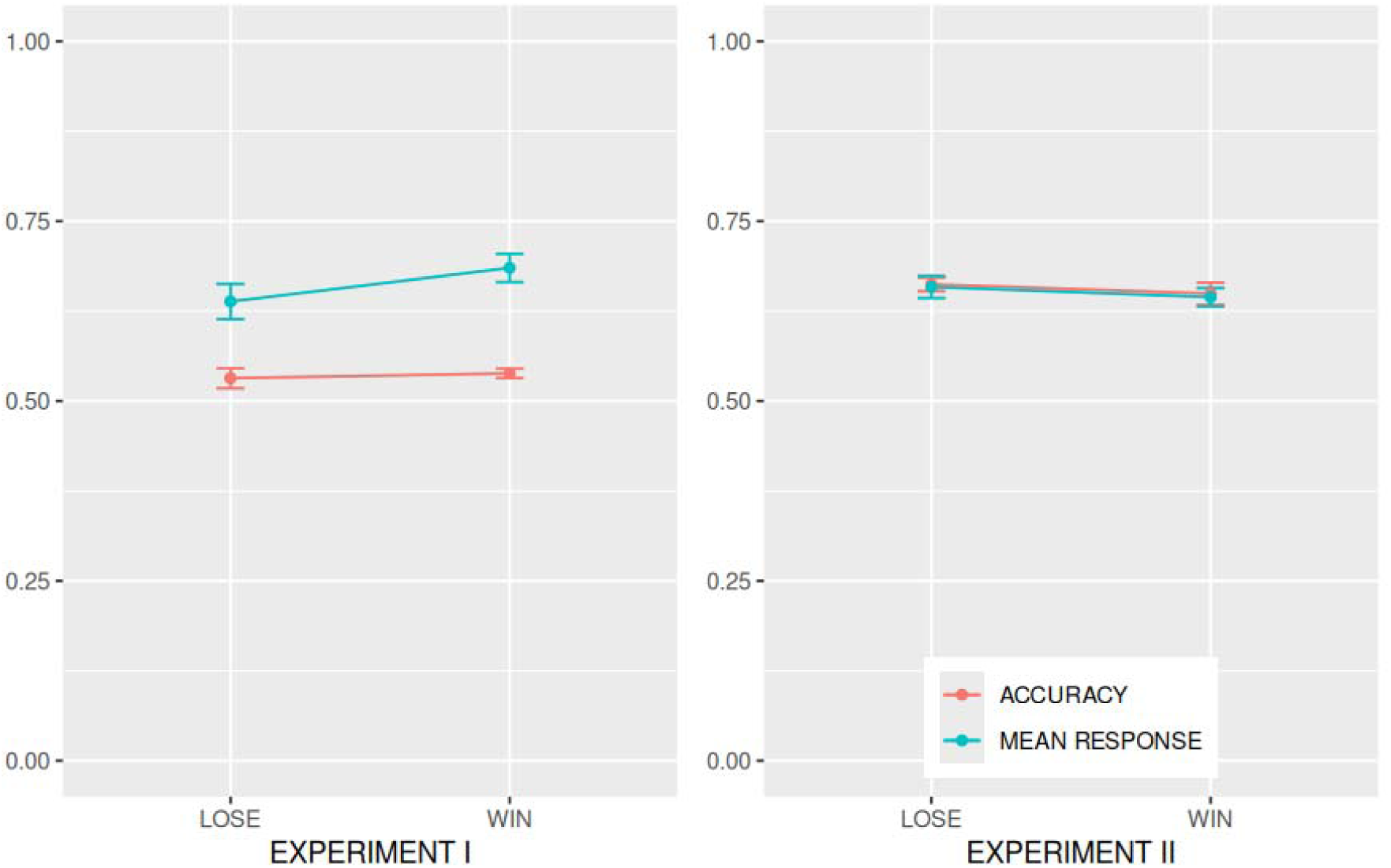
Accuracy and mean response for the two experimental groups. The “Win” group was rewarded 5 pence for each correct prediction, whereas the “Lose” group was punished with the loss of 5 pence for each incorrect prediction. Error bars are 95% confidence intervals.

As a simple visual inspection of Figure 13 shows, the higher mean accuracy achieved by participants in both groups of Experiment II compared with Experiment I strongly suggests that they underwent at least some degree of learning of the second—order Markov matrix’s temporal structure, which was absent from the zeroth—order input used in Experiment I. However, accuracy alone can only reveal this much about the participants’ decision—making behaviour. As we shall see, the toolkit proposed here is also capable of probing into the details of the putative learning process.

#### b. Cross—correlation function

The statistical analysis of the CCF was performed by a mixed—design 2 × 2 × 6 ANOVA, with Group (Win vs. Lose) as a between—subjects factor, Sequence (Output vs. Surrogate) and Lag (time lags from 0 to 5) as within—subjects factors. Figure 14 shows the cross—correlation functions for both the output sequences and their surrogate counterparts across time lags ranging from —5 to 5. Lags from —5 to —1 are included solely for visualisation purposes and were excluded from the statistical analyses, as they correspond to less meaningful correlations between output events and future input events. The Greenhouse—Geisser correction was applied because violations of sphericity were detected. The cross—correlation could reveal aspects of the participants’ decisional behaviour that the simpler measures of accuracy and mean response were unable to. Figure 14 reveals a clear asymmetry in the temporal profile of the CCF for both groups in both experiments, suggesting that the serial structure of the output sequence is influenced by past events in the input sequence (and, as expected, not the other way around).

**Figure 14.**
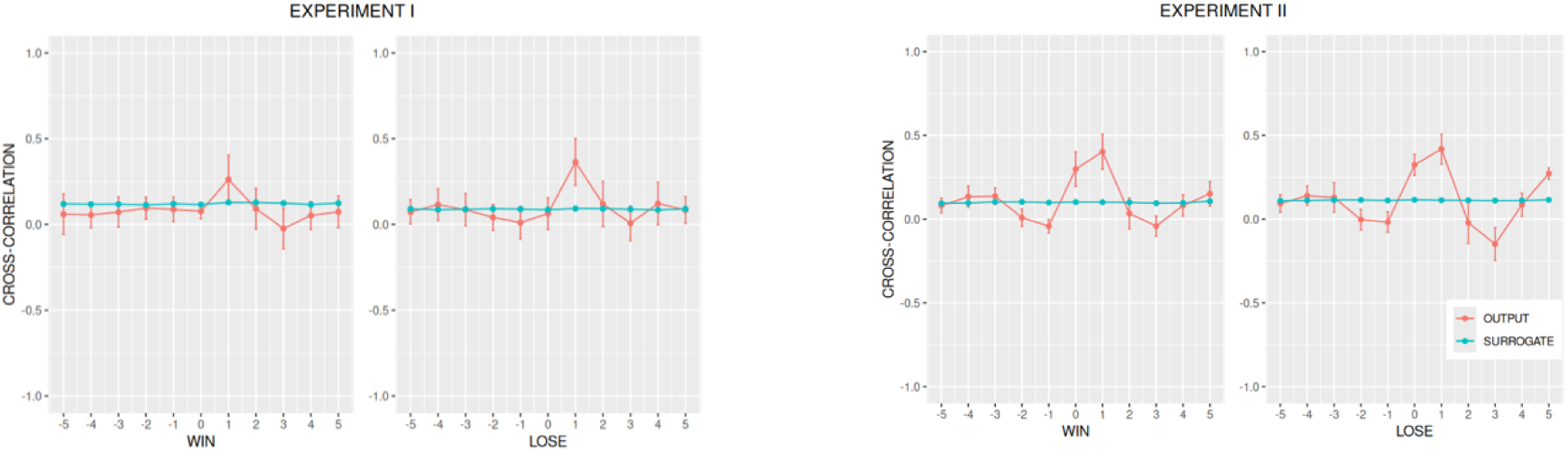
Cross—correlation of the output sequences and their surrogates as a function of the number of time intervals (lag). In red we can see the cross—correlation functions averaged across participants’ output sequences for both experimental groups (Win and Lose) in both experiments (Experiment I and Experiment II). A set of 100 surrogate sequences was obtained from each original output sequence by a random permutation of its elements. After calculating the mean cross—correlation function for each set of 100 surrogates, a grand mean was obtained by averaging the cross—correlation functions across the set of 50 functions, separately for each order and behaviour (teal lines). Error bars are 95% confidence intervals.

This influence becomes more evident when we find that a “significant” interaction between the factors Sequence and Lag (*F*(3.46, 34.58) = 12.340, *p* < 0.001, η_p_^2^ = 0.57 in Experiment I and *F*(2.24, 58.16) = 49.260, *p* < 0.001, η_p_^2^ = 0.65 in Experiment II) is mostly due to the peak at τ = 1 in the CCF of both Win and Lose groups in both experiments.

As explained earlier (see the discussion on the cross—correlation function in Simulation of Prototypical Behaviours — Results and Partial Discussion), a high peak in the CCF at τ = 1 strongly suggests that a win—stay/lose—shift (WSLS) strategy may substantially contribute to the overall decisional behaviour. However, no main effect (Exp I: *p* = 0.951; Exp II: *p* = 0.835) or interaction (Exp. I: *p* > 0.073, Exp. II: *p* > 0.122) involving the factor Group was strong enough to evidence a “statistically significant” difference between them.

#### c. Autocorrelation function

The statistical analysis of the ACF was performed by a mixed—design 2 × 2 × 5 ANOVA, with Group (Win vs. Lose) as a between—subjects factor, Sequence (Output vs. Surrogate) and Lag (time lags from 1 to 5) as within—subjects factors. The Greenhouse—Geisser correction was applied because violations of sphericity were detected. For Experiment I, this analysis revealed “significant” interactions between Group and Sequence (*F*(1, 10) = 8.040, *p* = 0.018, η_p_^2^ = 0.45) as well as between Lag and Sequence (*F*(1.93, 19.32) = 4.400, *p* = 0.028, η_p_^2^ = 0.32). For Experiment II, the interaction between the factors Group and Sequence was “not significant”, but the interaction between Lag and Sequence (*F*(2.33, 60.53) = 10.510, *p* < 0.001, η_p_^2^ = 0.29) was “significant”.

The “statistically significant” interaction between Group and Sequence in Experiment I indicates that the temporal correlation of the output sequences (averaged over the time lags) decays differently in the two groups, an effect that neither accuracy nor mean response was able to reveal. The interaction between Sequence and Lag also means that the temporal correlation of the output sequence (averaged over the two groups) decays with the lag in both experiments (since the ACF of the surrogate sequences does not change with the lag). These results are illustrated in Figure 15: the autocorrelation coefficients for the original output and surrogate sequences were qualitatively different from each other at τ = 1 in the Win group and at τ = 2 in the Lose group.

**Figure 15.**
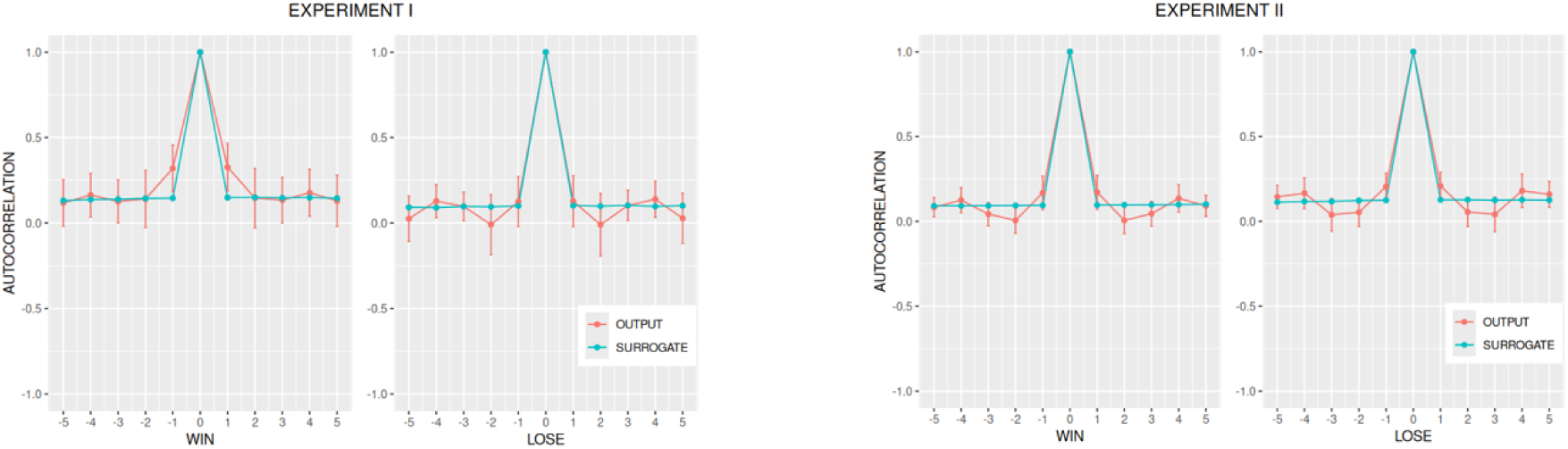
Autocorrelation of the output sequences and their surrogates as a function of the number of time intervals (lag). In red, we can see the autocorrelation functions averaged across participants’ output sequences for both experimental groups (Win and Lose) in both experiments (Experiment I and Experiment II). A set of 100 surrogate sequences was obtained from each original output sequence by a random permutation of its elements. After calculating the mean autocorrelation function for each set of 100 surrogates, a grand mean was obtained by averaging the autocorrelation functions across the set of 50 functions, separately for each order and behaviour (teal lines). The ACF values are not shown for lag τ = 0 since they are trivially identical to one. Error bars are 95% confidence intervals.

The interpretation of the image is straightforward: participants in the Win group showed a greater tendency than those in the Lose group to repeat their immediately previous choice (indicated by a positive coefficient on τ = 1); conversely, participants in the Lose group show a greater tendency, compared with those in the Win group, to choose an alternative opposite to the one chosen two steps earlier, as indicated by a negative coefficient on τ = 2. As we will see next, these interpretations are in line with the analysis of the shifting probabilities.

#### d. Shifting probabilities

Although a multivariate analysis of the shifting probabilities did not reveal a “significant” omnibus effect (Exp I: *Wilks’ lambda* = 0.573, *F*(2, 9) = 3.352, *p* = 0.0816, η_p_^2^ = 0.43; Exp. II: *Wilks’ lambda* = 0.983, *F*(2, 25) = 0.211, *p* = 0.811, η_p_^2^ = 0.02), Figure 16 shows that the two dependent variables, *P*(shift|win) and *P*(shift|lose), tend to exhibit a higher value in the participants of the Lose group when compared to those in the Win group in Experiment I, but not in Experiment II. In fact, in Experiment I, a univariate ANOVA (which must be interpreted with caution following a “nonsignificant” MANOVA) was able to show a marginally “significant” difference between the two groups for *P*(shift|lose) (*F*(1, 10) = 7.113, *p* = 0.023).

**Figure 16.**
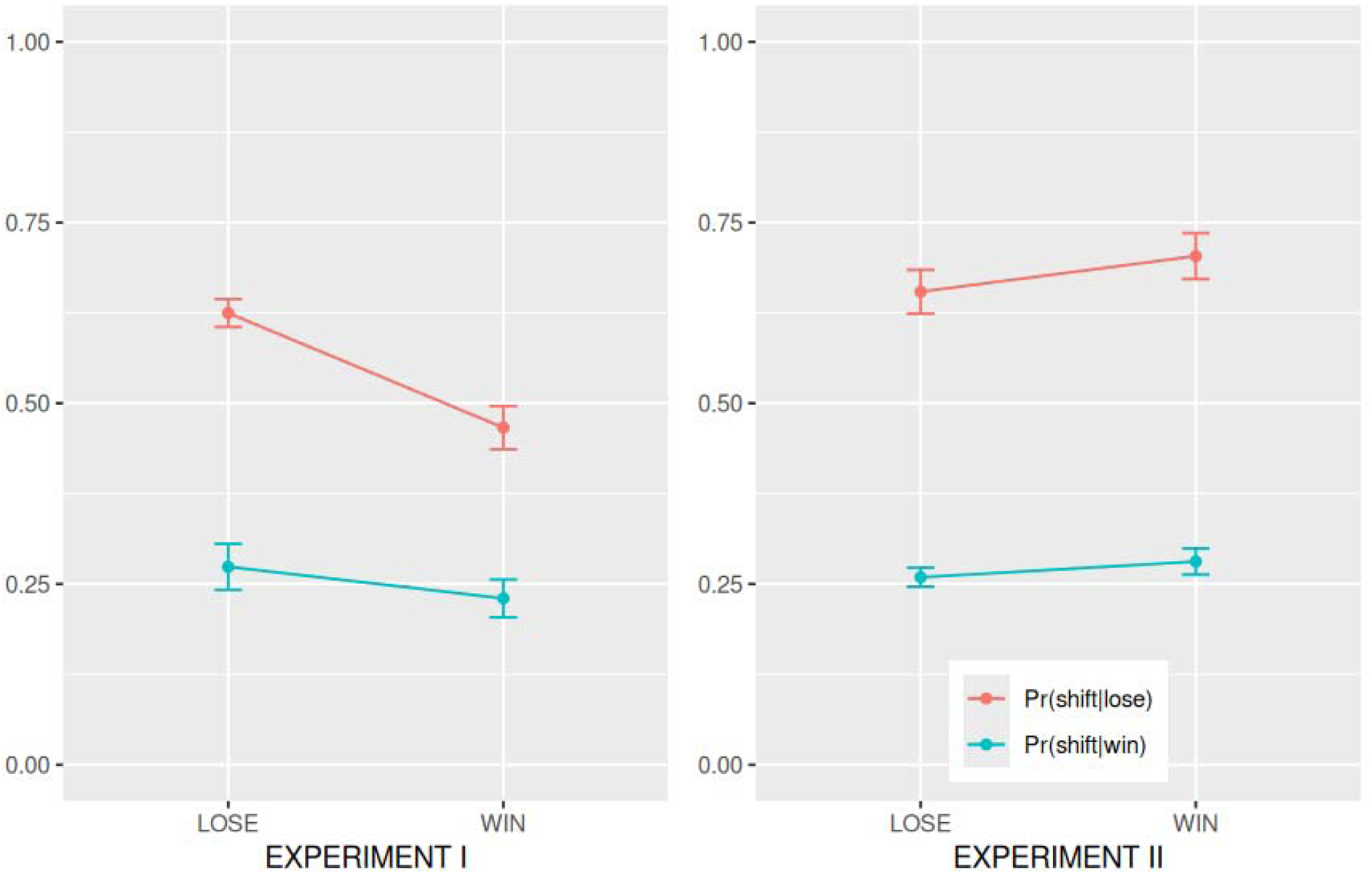
Unconditional probability of shifting, P(shift), and conditional probabilities of shifting given that the participant’s previous prediction was correct, P(shift|win), or incorrect, P(shift|lose), for both experimental groups (Win and Lose) and both experiments (Experiment I and Experiment II). Error bars are 95% confidence intervals.

The effect of differential payoffs on the shifting probabilities, even though not “statistically significant”, is qualitatively consistent with those revealed by the analysis of the autocorrelation and cross—correlation functions. Altogether, these analyses point to a more exploratory behaviour of the Lose group and a more exploitative behaviour of the Win group in Experiment I. This behavioural distinction is supported by (i) a “significant” positive ACF(1) found in the Win group and absent in the Lose group, (ii) a “significant” negative ACF(2) found in the Lose group (and absent in the Win group), (iii) a higher — and “statistically significant” — positive peak in CCF(1) for the Lose group in comparison to the Win group, and (iv) a qualitative inclination towards higher values for all three shifting probability measures in the Lose group when compared with the Win group.

A qualitative analysis that lends further support to the conclusions above is provided by the shifting diagram, in which a scatterplot of *P*(shift|win) against *P*(shift|lose) is depicted for both experimental groups (Figure 17). According to our previous description of the shifting diagram (see Figure 1), we can notice that, in Experiment I, all participants in the Lose group are confined to the win—stay/lose—shift quadrant, whereas at least half of the participants in the Win group clearly fall within the win—stay/lose—stay quadrant.

**Figure 17.**
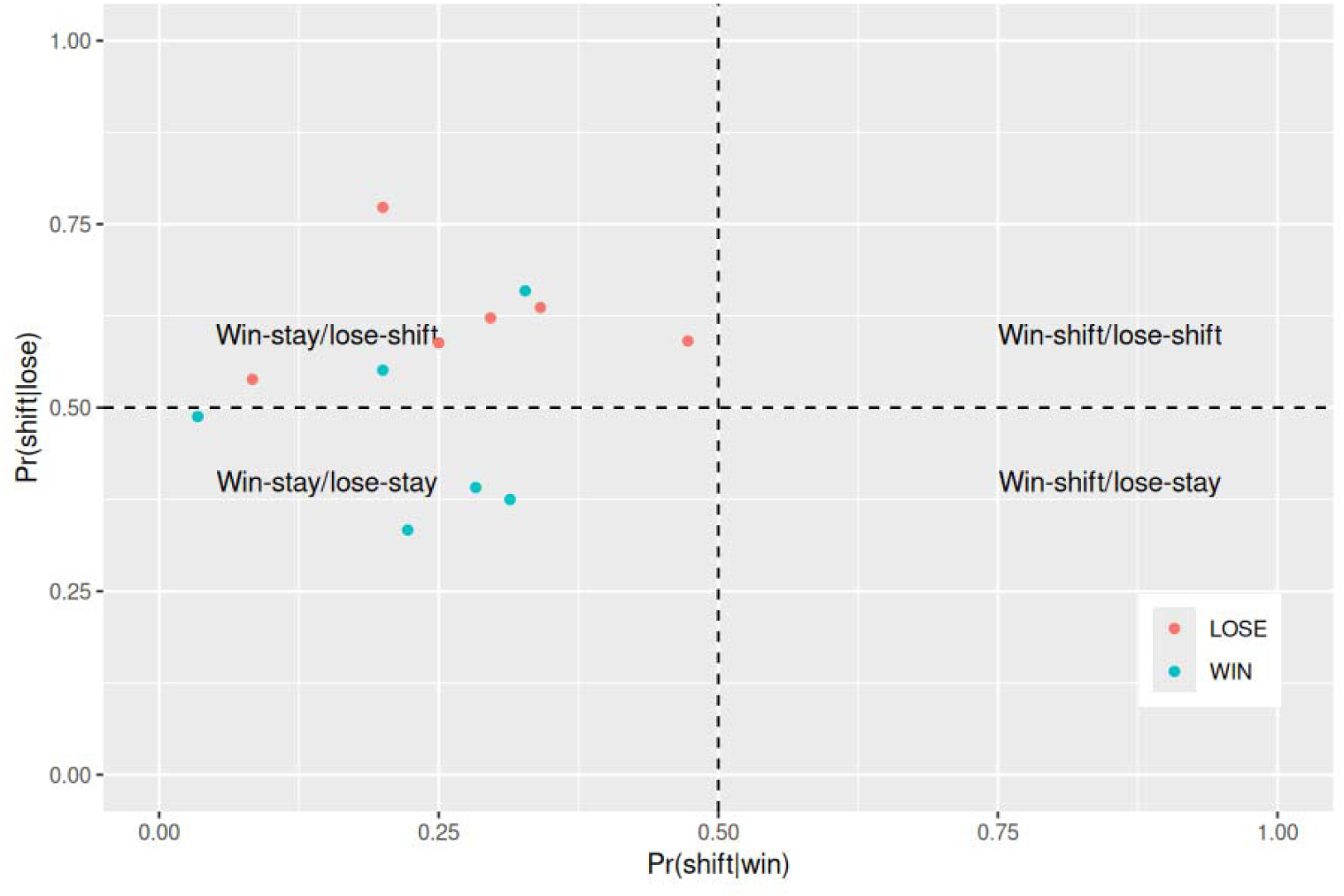
Shifting diagram showing a scatterplot of P(shift|win) against P(shift|lose) for both experimental groups (Win and Lose) in Experiment I. Whereas all participants in the Lose group (red dots) occupy the win—stay/lose—shift quadrant (top—left), four out of six participants in the Win group (teal dots) occupy the win—stay/lose—stay quadrant (bottom—left).

In Experiment II, the differences in shifting probabilities between the Win and Lose groups are not “statistically significant”, and most participants in both groups fall within the win—stay/lose—shift quadrant (*Figure 18*; see e. Shifting probabilities).

**Figure 18.**
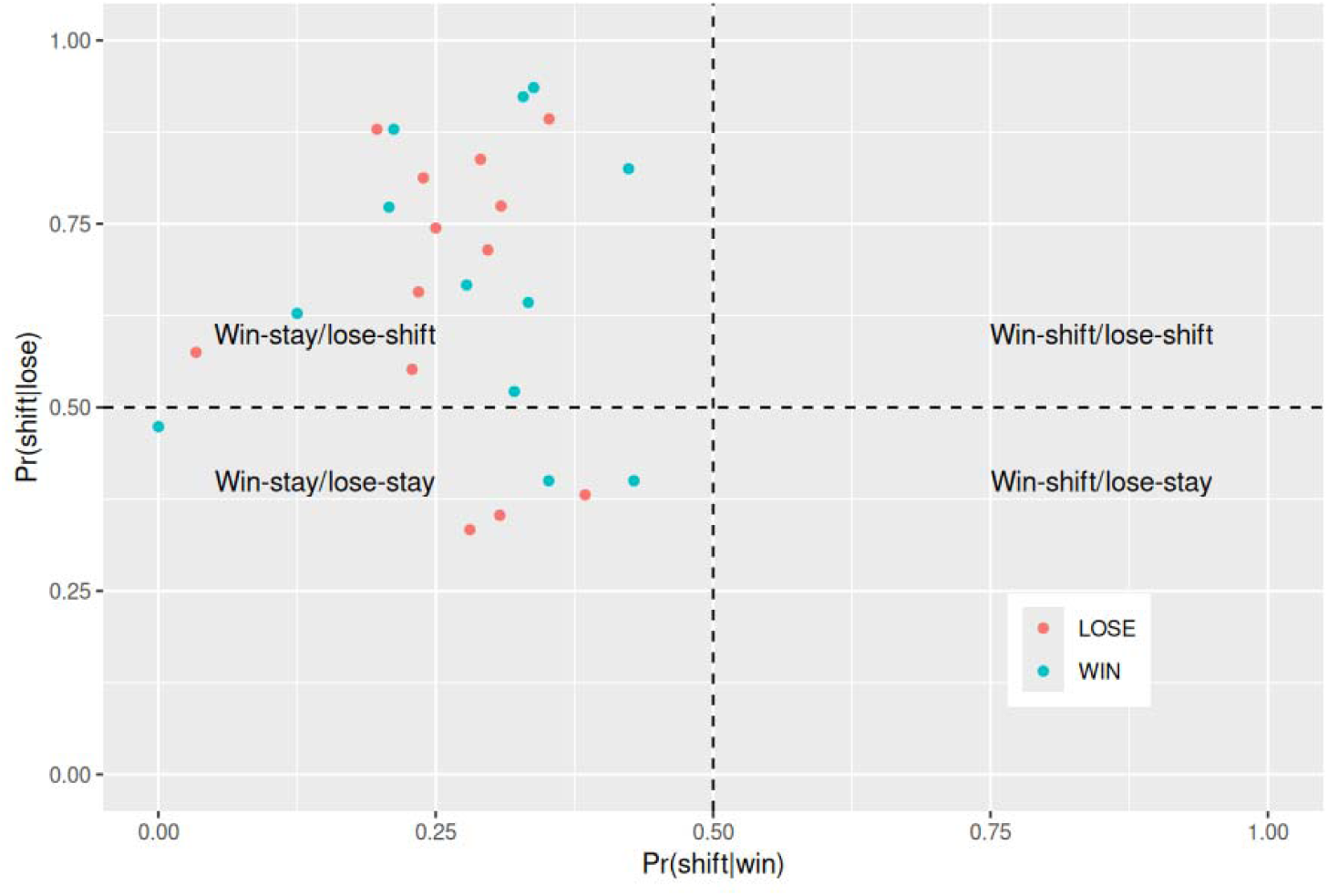
Shifting diagram showing a scatterplot of P(shift|win) against P(shift|lose) for both experimental groups (Win and Lose) in Experiment II. Almost all volunteers in the Lose group (red dots) and Win group (teal dots) occupy the win—stay/lose—shift quadrant (top—left), with six other volunteers (three from each group) lying in the win—stay/lose—stay quadrant (bottom—left).

#### e. Markov reconstruction

The Markov reconstruction expresses the participants’ asymptotic behaviour for each experiment. Table VI shows that both groups (Win and Lose) in Experiment I were able to learn the most frequent element of the sequence, since in both cases the probabilities of choosing the most frequent event (+1) were “significantly” higher than 0.5, with a 95% confidence interval of [0.62, 0.75] and [0.56, 0.72], respectively.

**Table VI.** Zeroth—order Markov matrices reconstructed from the output sequences generated by Experiment I participants in response to an M0 input sequence. The value of each element in the matrices is expressed by a 95% confidence interval calculated from the set of 12 volunteers.

| EXPERIMENT I — WIN |  | EXPERIMENT I — LOSE |  |
| --- | --- | --- | --- |
| $y_i$ | | $y_i$ | |
| –1 | +1 | –1 | +1 |
| [0.25, 0.38] | [0.62, 0.75] | [0.28, 0.44] | [0.56, 0.72] |

In Experiment II, participants in both groups (Win and Lose) generated Markovian sequences that, at least partially, mimicked the Markov matrix that generated the input sequence, as shown in Table VII. This is also supported by comparing the results for both groups with those of their respective surrogates (Figure 19). The Markov matrix elements of the surrogates approximate the transition probabilities of a zeroth—order Markov chain by destroying the temporal correlations in the output sequences. This result lends support to the conclusion that participants in both groups were able to learn a significant portion of the information contained in the second—order input sequence.

**Figure 19.**
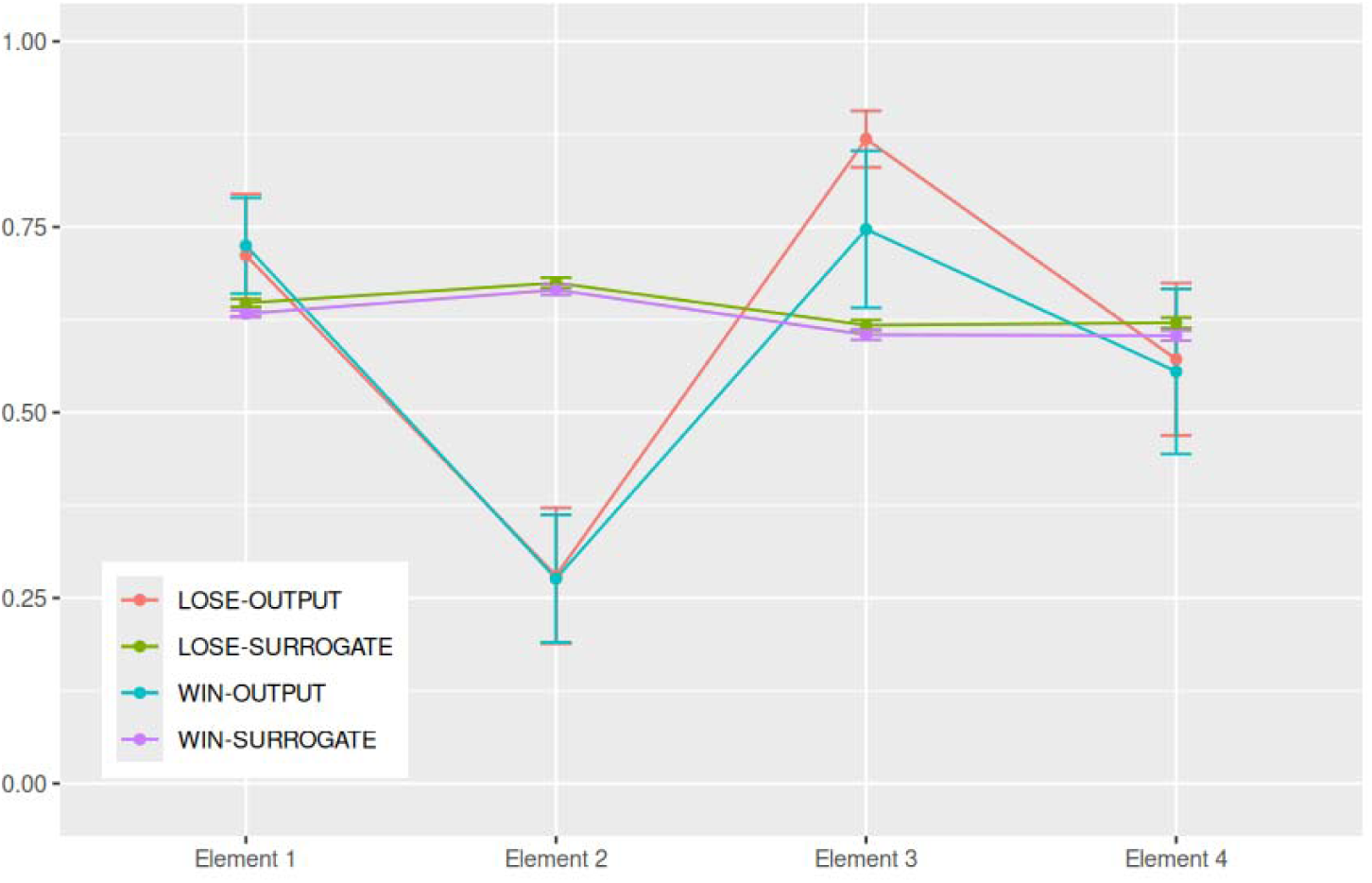
Graph showing the four elements of the second—order Markov matrices reconstructed from the output sequences generated by Experiment II participants in response to an M2 input sequence. Error bars are ± 1 standard deviation.

**Table VII.**
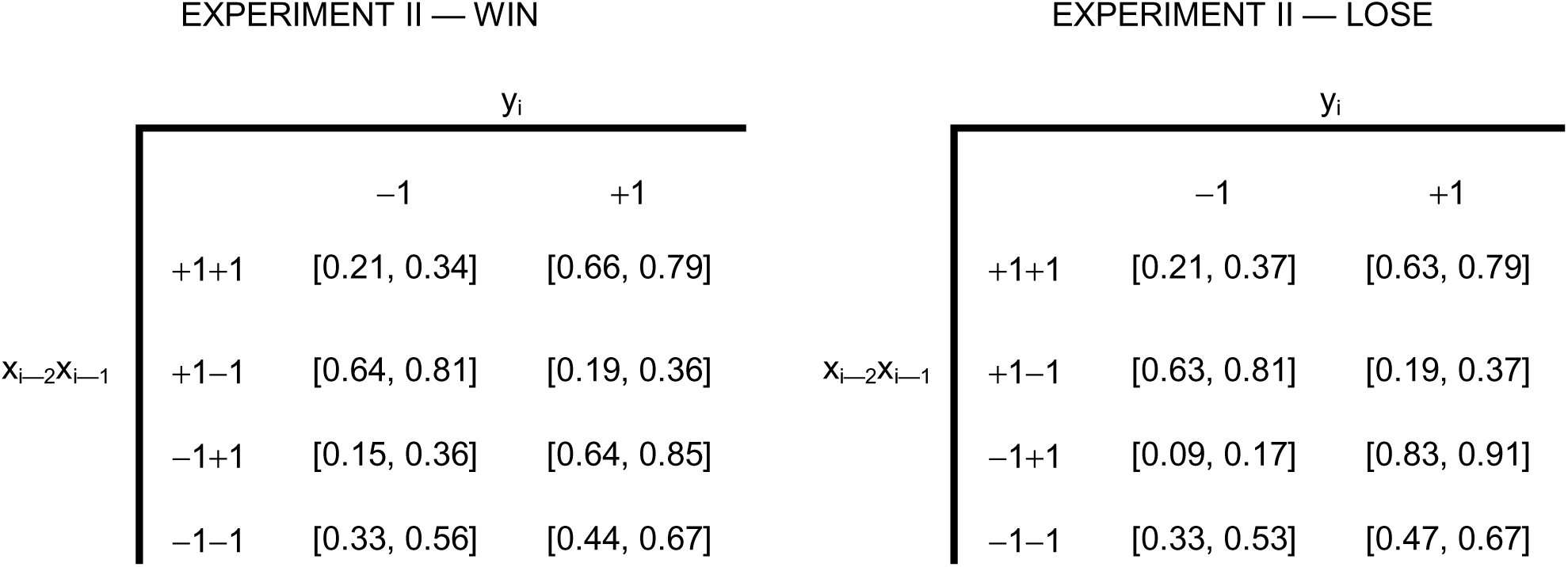
Second—order Markov matrices reconstructed from the output sequences generated by Experiment II participants in response to an M2 input sequence. The value of each element in the matrices is expressed by a 95% confidence interval calculated from the set of 28 volunteers.

#### f. Partial discussion

In conclusion, in Experiment I, differential payoffs (rewards or punishments) produced a clear, distinctive bias in the decisional strategy adopted by the two experimental groups when predicting a sequence with a more frequent element. Despite being submitted to the same binary choice task, volunteers in the Win group tended towards a more exploitative behaviour, whereas those in the Lose group adopted a more exploratory strategy.

For Experiment II, the volunteers were able to extract most of the information from the task and partially learn the sequence structure, which is evidenced mainly by the Markov reconstruction. However, these conclusions were only possible through the analysis of multiple complementary measures that jointly extract valuable information from a probability learning task involving binary sequences — information that would have remained hidden had the analysis relied solely on mean response and accuracy (which, in Experiment II, could only indicate the likely presence of temporal structure learning).

## General Discussion

The present study aimed to introduce and validate a computational toolkit for the analysis of repeated binary choice tasks, leveraging techniques that go beyond conventional accuracy and response frequency metrics, which obscure the rich temporal structure embedded in sequential behaviour. By integrating methods such as autocorrelation, cross—correlation, shifting probabilities, graphical analysis, and the reconstruction of Markov transition matrices, we were able to extract structural elements from both simulated and empirical binary sequences that would otherwise remain inaccessible.

Taken together, the set of metrics introduced in this toolkit provides a complementary and multidimensional characterisation of binary choice behaviour, capturing both aggregate tendencies and temporal structure. Mean response offers a simple estimate of overall bias toward one option, while accuracy reflects performance relative to the sequence to be predicted. However, as demonstrated here, these measures alone may be insufficient and, in some cases, potentially misleading when interpreted in isolation. We note that the mean response and accuracy can in principle detect group differences in many contexts, although they did not reveal “statistically significant” differences in our empirical data. More importantly, the main advantage of the toolkit lies in uncovering how participants behave in finer detail within each group. The cross—correlation quantifies the alignment between the sequence to be predicted (input sequence) and the sequence generated by the participant (output sequence), across time lags, revealing whether and how past events influence current decisions, whereas the autocorrelation captures internal dependencies within the response sequence itself, indicating persistence or alternation tendencies. Shifting probabilities provide an intuitive behavioural summary of local decision rules, specifically, sensitivity to previous outcomes (e.g., Win—Stay/Lose—Shift), and allow for direct mapping onto well—known heuristics. Finally, Markov reconstruction offers a more formal and model—based perspective by estimating the conditional structure governing responses, thereby enabling the identification of the temporal dependencies underlying behaviour. Importantly, although some degree of redundancy exists within these measures, they operate at different levels of description and, when used jointly, permit a richer and more precise inference of the strategies and dynamics that shape sequential decision—making.

Additionally, by introducing Markov binary chains of varying orders, it becomes possible to generate input sequences (i.e., the sequences to be predicted) with controllable levels of serial complexity, thereby more closely approximating real—world environments, in which events are rarely independent. This enables us to investigate more thoroughly the probability learning capacity of agents — both simulated models and human or nonhuman animals — when challenged to predict sequences of varying complexity.

The application of these methods to prototypical behaviours provided a useful interpretative bridge between abstract mathematical measures and psychologically meaningful strategies, as demonstrated in the simulations (Simulation of Prototypical Behaviours — Results and Partial Discussion). For instance, the identification of a peak at lag τ = 1 in the cross—correlation function offers a clear diagnostic marker of WSLS—like behaviour (mathematically indistinguishable from simply replicating the previous event), while oscillatory patterns in higher—order contexts reveal sensitivity to more complex temporal dependencies. Similarly, shifting probabilities provide an intuitive yet powerful way to map behaviour onto well—established decision—making heuristics, capturing the balance between exploitation and exploration.

When applied to real empirical data, our analysis toolkit successfully uncovered strategy differences between groups of participants subjected to differential reinforcement regimes — reward (Win) versus punishment (Lose) — and sequence complexities — zeroth— versus second—order Markov chains. While simple measures such as mean response and accuracy failed to reveal “significant” differences between these groups, more refined analyses revealed distinct serial structures in the participants’ choice patterns. Specifically, for Experiment I, which employed a zeroth—order Markov chain as the input sequence, participants in the Win group displayed stronger exploitative tendencies, evidenced by higher autocorrelation at lag 1, lower shift probabilities, and a clustering in the win—stay/lose—stay quadrant of the shifting diagram. Conversely, the Lose group exhibited more exploratory behaviour, reflected in higher shift probabilities, a “statistically significant” negative autocorrelation at lag 2, and a clear placement within the win—stay/lose—shift quadrant. Such a pattern aligns with the findings of Tversky & Kahneman (1981), according to which choices involving gains tend to be risk—averse, whereas choices involving losses promote risk—taking. It is also consistent with the results reported by Bereby—Meyer & Erev (1998), which indicate that learning may proceed more rapidly when reinforcement is based on losses.

It is worth noting that the potential of the present techniques to reveal differences between the two groups persisted even under conditions of relatively low statistical power (i.e., small sample sizes and short binary sequences, which, in empirical studies, is sometimes imposed by technical constraints). The lack of adequate statistical power likely contributed to the failure of simpler analyses — based on accuracy and mean responses — to yield “statistically significant” results, while not substantially compromising the performance of at least some of the techniques in the present analytical toolkit.

For Experiment II, which employed a second—order Markov chain as the input sequence, analyses based solely on mean response and accuracy also failed to reveal “statistically significant” differences in participants’ choices. However, accuracy itself was sufficient to indicate that participants had learned at least some aspects of the temporal structure of the M2 sequence, as performance exceeded what would be expected from reliance on simple event frequencies. What accuracy could not reveal was which aspects of the temporal structure were learned. The complementary analyses introduced in the present toolkit provided a more detailed characterisation of participants’ behaviour. The cross—correlation function provided initial evidence that participants’ responses were not independent of the input sequence, revealing lag—dependent structure consistent with sensitivity to preceding events. The autocorrelation function further supported this view by showing non—trivial internal dependencies in the response sequences, inconsistent with purely stochastic responding. Most importantly, a second—order Markov chain was reconstructed from the participants’ data, revealing that decisional behaviour was strongly guided by preceding elements of the input sequence and providing clear evidence of learning.

Taken together, our findings strongly support the notion that subtle yet measurable aspects of decision—making strategies or learning capabilities might be easily overlooked in small samples when relying exclusively on traditional behavioural metrics.

In summary, the proposed toolkit expands the analytical repertoire available for studying repeated binary choice tasks, enabling a more nuanced and structurally informed characterisation of behaviour. By revealing patterns that remain undetected by traditional measures, it opens new avenues for investigating the cognitive and neural mechanisms underlying probabilistic learning and decision—making, with potential applications spanning domains ranging from basic research to clinical assessment.

### 1. Limitations and Future Directions

While the present study introduces a comprehensive computational toolkit for analysing repeated binary choice behaviour, several limitations should be acknowledged, both in the methodological scope of the framework and in its current empirical validation.

A first limitation concerns the reliance on simulated prototypical behaviours as a primary means of validating the analytical methods. Although these simulations are useful for establishing clear mappings between known strategies and their statistical signatures, they necessarily simplify the complexity of real behaviours. In practice, participants are unlikely to follow pure strategies such as strict probability matching or ideal prediction. Instead, behaviour is often noisy, non—stationary, and composed of mixtures of strategies that evolve over time. As a result, the interpretability of the proposed measures may be reduced when applied to empirical data where multiple latent processes interact.

A second limitation of this study is that the results presented here are largely contingent on the specific input sequences used in the simulations and cannot be generalised to other input sequences except for a small number of invariants. However, we also provide a detailed description of the procedures as well as the code used to generate these results, enabling other researchers to readily explore and evaluate sequences produced by any Markov processes.

A third limitation relates to the assumption of stationarity in both input and output sequences. Many of the analytical tools employed here — such as autocorrelation, cross—correlation, and Markov reconstruction — implicitly assume that the statistical properties of the sequences remain constant over time. However, learning is inherently a dynamic process: participants may gradually acquire knowledge about the structure of the sequence, change strategies, or adapt to changes in reward contingencies. By discarding early trials and analysing only approximately stationary segments, the present approach may overlook important aspects of learning dynamics. Future developments should therefore aim to incorporate analyses of participants’ learning kinetics.

Looking forward, future work could extend the analytical tools proposed here by incorporating additional analytical tools — such as entropy rate, hidden Markov models, or other measures of dissimilarity between matrices — to better capture the dynamics of the strategies underlying binary choice behaviours. Also, by exploring more complex probabilistic learning scenarios, such as those involving two or more agents interacting on the same prediction task, the prediction of more than two alternatives or the introduction of real—time feedback dynamically shaping the stimulus sequence, the scope of the present toolkit could be largely expanded, offering an approach relevant to both basic and clinical research.

In conclusion, despite these limitations, the proposed analytical methods offer a valuable and versatile approach to the study of binary choice behaviour. By enabling a deeper structural analysis of sequential choice behaviour, it opens new possibilities for understanding the cognitive strategies that govern human and animal choices under uncertainty.

## Supplementary Information

The online version contains supplementary material available at https://github.com/bernard17costa/paper_computational_toolkit

## Acknowledgement

The authors would like to express their gratitude to the late Prof. Glyn Humphreys, formerly Watts Professor of Experimental Psychology and principal investigator of the CNN Lab at Oxford University, who generously hosted one of the authors (MB) as a visiting researcher and provided valuable material and intellectual support to this work.

## Author Contributions

Bernard Costa: Data collection and curation, Formal analysis, Methodology, Writing — original draft, Writing — review & editing.

Marcus Baldo: Conceptualisation, Data collection and curation, Formal analysis, Methodology, Writing — original draft, Writing — review & editing, Funding acquisition, Supervision.

Carolina F. da Silva: Methodology, Writing — original draft, Writing — review & editing, Supervision.

## Funding

This research was funded by a (CAPES) fellowship to BC (88887.748837/2022—00) and a FAPESP fellowship to MB (2013/13352—2).

## Availability of data, codes and materials

Data and code used for the analyses are available on the https://github.com/bernard17costa/paper_computational_toolkit

## Declarations

### Conflicts of interest

The authors declare no conflicts of interest.

### Ethics approval

The experimental procedures were approved by the Ethics Committee of the Department of Experimental Psychology, University of Oxford, and the Ethics Committee of the Institute of Biomedical Sciences, University of São Paulo.

### Consent to participate

Informed consent was obtained from all individuals involved in the study or, in the case of underage individuals, from their parents or legal guardians. Individuals’ data were anonymised by replacing their names with numbers in the data set.

### Generative AI statement

During the preparation of this work, the authors used generative AI and AI—assisted technologies (GPT—5.1, ChatGPT, OpenAI; and Claude Sonnet 4.6, Anthropic) solely to improve the language and readability of the manuscript — specifically grammar, spelling, punctuation, agreement, and stylistic clarity. The tools were not used for study design, creation of scientific content, data extraction, figure or table generation, data analyses, or interpretation. After using these tools, all AI—assisted suggestions were manually reviewed, edited, and approved by the authors, who take full responsibility for the accuracy and integrity of the final manuscript.

### Open practices statement

None of the reported analyses was preregistered. A previous version of this work was deposited as a preprint in bioRxiv prior to peer review. The preprint is available at https://doi.org/10.64898/2026.01.21.700416, and includes revisions and updates made after the initial preprint release.

## Notes

https://github.com/bernard17costa/paper_computational_toolkit

### Competing Interest Statement

The authors have declared no competing interest.

### Summary of Updates

Updates throughout the text, especially in the part describing the experiments with human participants.

